# Structural Basis of Substrate Selectivity and Catalysis in the Mycobacterial Long-Chain Acyl-CoA Carboxylase

**DOI:** 10.64898/2026.02.06.704326

**Authors:** Ajit Yadav, Nicole Rizzetto, Bogdan I. Florea, Sebastian Geibel

**Affiliations:** Leiden Institute of Chemistry, Leiden University, Leiden, Netherlands

## Abstract

Long-chain acyl-CoA carboxylase (LCC) is an essential enzyme complex in mycobacteria that generates acyl-CoA precursors for mycolic acid and complex lipid biosynthesis, yet its architecture and mechanism of substrate selection have remained unclear. Here we determine pre- and post-reaction states of the endogenous 868-kDa LCC complex from *Mycobacterium smegmatis* by cryo-electron microscopy at 2.1-3.7 Å resolution. These structures visualize ATP-dependent redistribution of the biotin carboxyl carrier protein. LCC assembles into an asymmetric 8:2:4:2 organization of AccA3, AccD4, AccD5, and AccE5, with two biotin carboxylase modules flexibly tethered to a heterohexameric carboxyltransferase core. We define the structural basis of substrate selectivity within the CT core: AccD5 selectively binds the short-chain substrate C3-CoA, whereas AccD4 accommodates the long-chain substrate C16-CoA.

In addition, we resolve AccD5-centered assemblies that associate with biotin carboxylase modules yet lack AccD4, providing structural evidence that distinct carboxyltransferase cores can engage shared modules to generate alternative holoenzyme architectures. Together, these findings define LCC and AccD5-centered assemblies as elements of a combinatorial acyl-CoA carboxylase platform and establish the structural principles governing assembly-specific function in mycobacteria.

## Introduction

A hallmark of mycobacteria, including major pathogens such as *Mycobacterium tuberculosis* and *Mycobacterium abscessus* as well as environmental species such as *Mycobacterium smegmatis*, is a highly impermeable, lipid rich outer membrane known as the mycomembrane^1,2,3^. This barrier underlies intrinsic antibiotic tolerance, environmental resilience, and, in *M. tuberculosis*, long term persistence within the host^4,5^. Maintenance of mycomembrane integrity depends on the biosynthesis of mycolic acids, long α-alkyl, β- hydroxy fatty acids whose extreme chain length and structural complexity require specialized metabolic machinery^6,7,8^. In addition to mycolic acids, the mycomembrane contains diverse complex lipids that contribute to envelope architecture and permeability, and whose biosynthesis depends on the availability of acyl-CoA precursors^7^.

In *M. tuberculosis*, the long-chain acyl-CoA carboxylase (LCC) complex catalyzes Cα- carboxylation reactions that contribute to methylmalonyl-CoA production for multimethyl- branched cell envelope lipids, such as phthiocerol dimycocerosates, and generate the α-branch precursors required for mycolic acid biosynthesis ^9,10,11^. Through these reactions, LCC occupies a central metabolic position linking carbon metabolism to cell envelope lipid production. Catalysis proceeds via the canonical two step, biotin dependent mechanism, in which ATP driven carboxylation of biotin on the biotin carboxyl carrier protein by the biotin carboxylase activity is followed by carboxyl transfer to an acyl CoA substrate at the carboxyltransferase active site^12,13^.

Unlike organisms such as *Escherichia coli* or humans, *M. tuberculosis* encodes an expanded repertoire of acyl-CoA carboxylase subunits, comprising three biotin carboxylases (AccA1- AccA3) and six carboxyltransferases (AccD1-AccD6)^14^. Genetic and biochemical studies have linked specific α (biotin carboxylase) and β (carboxyltransferase) subunits to distinct metabolic functions via defined α/β pairings, including AccA3-AccD6 in acetyl-CoA carboxylation for fatty-acid biosynthesis^15^, AccA1-AccD1 in leucine degradation^16^, and AccA3-AccD5- dependent short-chain acyl-CoA carboxylation activities producing malonyl- and methylmalonyl-CoA (sometimes referred to as ACCase 5)^17–19^. Collectively, these assignments have largely relied on in vitro reconstitution or pairwise interaction assays, and the in vivo organization, stoichiometry, and modularity of mycobacterial acyl-CoA carboxylase assemblies remain poorly defined.

LCC is thought to assemble the biotin carboxylase AccA3, two carboxyltransferases AccD4 and AccD5, and the essential ε subunit AccE5 into a multifunctional enzyme complex^10^. Within this complex, long chain acyl CoAs ranging from C16 to C26 are processed by AccD4^18,20^, whereas short chain substrates, including acetyl-CoA and propionyl-CoA, are processed by AccD5^11,17,20^. Despite its central role in mycobacterial physiology, key properties of the endogenous LCC, including subunit stoichiometry, overall architecture, conformational dynamics, biotin carboxyl carrier protein (BCCP) trafficking, and the molecular basis of dual chain length specificity, have remained unresolved. Structural insight has so far been limited to isolated components, including an AccA3 dimer and an AccD5 hexamer ^11,21^. The organization of these subunits within intact assemblies is unknown. To address these gaps, we determined high-resolution cryo-EM structures of endogenous LCC across multiple catalytic states and defined the structural organization of AccD5-centered assemblies using single-particle cryo-electron microscopy.

## Results

### Serendipitous overproduction of endogenous LCC in *M. smegmatis* enables purification of the native complex

LCC was unexpectedly detected during StrepTactin pulldown experiments from *M. smegmatis* carrying an empty pMyNT plasmid (Fig. 1a). Owing to the covalently biotinylated AccA3 subunit, native LCC was purified directly from *M. smegmatis* lysate by StrepTactin affinity chromatography followed by size-exclusion chromatography. Acetamide induction markedly increased LCC yield, whereas only low amounts were obtained from uninduced cells. Cryo-EM particle classification revealed minor populations corresponding to additional carboxylase assemblies that co-purified with LCC.

**Figure 1.**
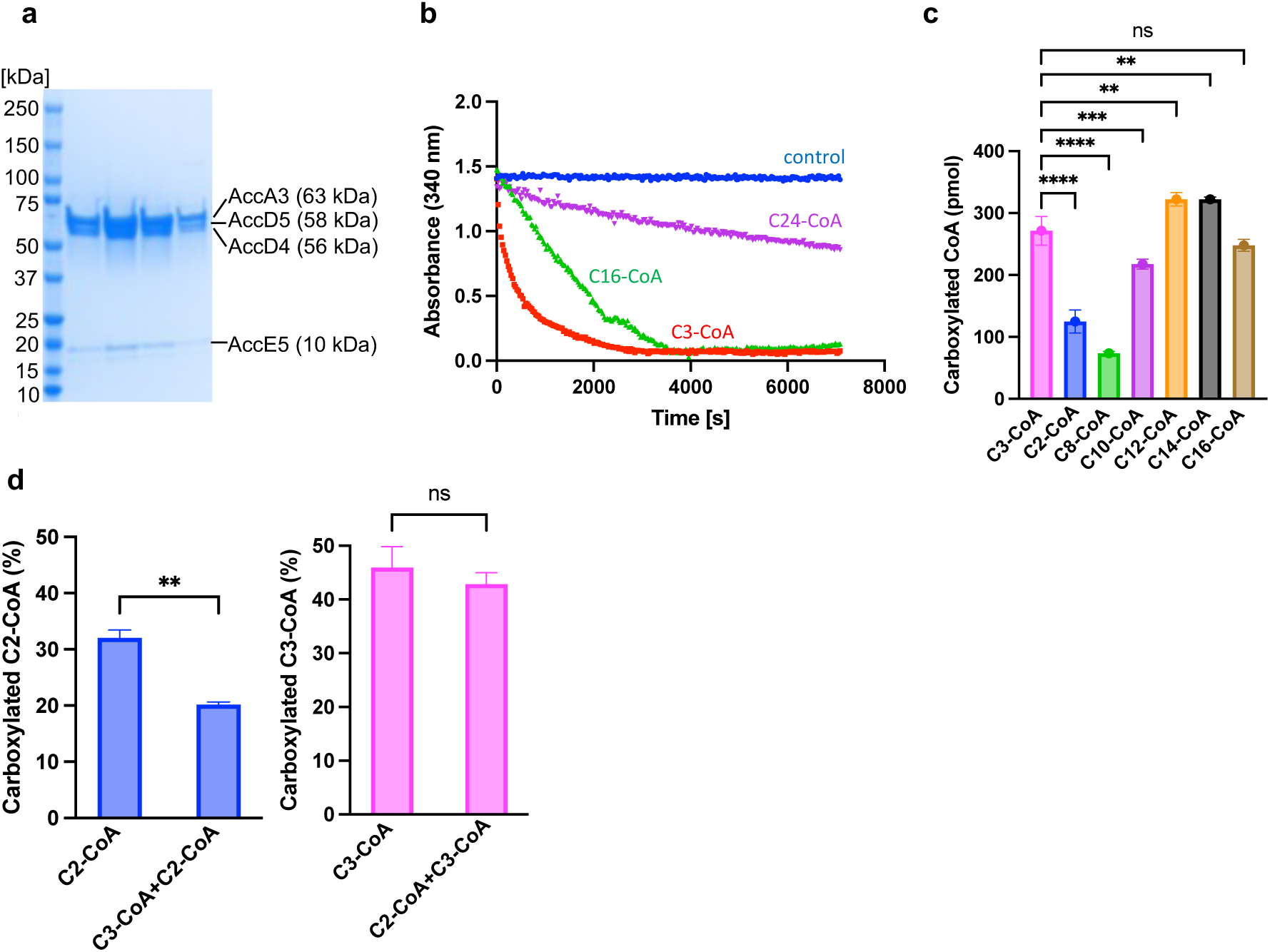
Purification and biochemical characterization of an endogenous acyl-CoA carboxylase preparation enriched in LCC. **a,** SDS-PAGE analysis of the endogenous acyl-CoA carboxylase preparation purified from *Mycobacterium smegmatis*, dominated by the long-chain acyl-CoA carboxylase (LCC) complex. **b,** Substrate-stimulated ATPase activity of the purified preparation measured in the presence of propionyl-CoA (C3-CoA), palmitoyl-CoA (C16-CoA), or lignoceroyl-CoA (C24-CoA). **c,** Quantification of carboxylated acyl-CoA species (C2-, C3-, C8-, C10-, C12-, C14-, and C16-CoA) detected by IP-RP-HPLC-ESI-TQMS/MS. **d,** Substrate-competition assay measuring carboxylation of C2-CoA and C3-CoA in the presence of the competing substrate. Statistical significance in **c** was assessed by one-way ANOVA (P < 0.0001) followed by Dunnett’s multiple-comparisons test (reference, C3-CoA; ****P < 0.0001, ***P < 0.001, **P < 0.01; ns, not significant). Welch’s unpaired two-tailed *t*-test was used in **d** (**P < 0.01; ns, not significant). Data are mean ± s.d. (n = 3).

### Endogenous LCC is catalytically active

Biochemical assays were performed on the endogenous acyl-CoA carboxylase preparation purified from *M. smegmatis*, which contains LCC together with co-purifying minor assemblies. The purified material exhibited activity in both half-reactions of the carboxylation cycle. ATPase activity was highest in the presence of propionyl-CoA (C3) and progressively lower with the long-chain substrates palmitoyl-CoA (C16) and lignoceroyl-CoA (C24) (Fig. 1b). LC-MS analysis revealed a broad but discontinuous substrate spectrum, with robust carboxylation of short-chain (C2, C3) and long-chain (C10-C16) acyl-CoAs (Fig. 1c; Supplementary Fig. 1). Activity toward the medium-chain substrate hexanoyl-CoA (C6) was detectable but below the limit of quantification, while octanoyl-CoA (C8) showed only minimal activity. Carboxylation of substrates longer than C16 was detectable but could not be quantified due to strong retention on the LC column (Supplementary Fig. 1-2). The observed substrate specificity is consistent with that reported for reconstituted *M. tuberculosis* LCC ^10^.

Carboxylation of long-chain acyl-CoA substrates requires AccD4 and therefore specifically reports on LCC activity, whereas short-chain activity (C2, C3) may additionally reflect contributions from the AccA3-AccD5-AccE5 (ACCase 5) assembly. In *M. smegmatis*, AccD5 supplies both malonyl-CoA^20^ and methylmalonyl-CoA^11^ via carboxylation of acetyl-CoA and propionyl-CoA, respectively, but the relative preference of *M. smegmatis* LCC for acetyl-CoA versus propionyl-CoA has not been defined. To probe substrate preference, we performed LC-MS-based competition assays, which showed that carboxylation of acetyl-CoA (C2) was significantly reduced in the presence of propionyl-CoA (C3), whereas carboxylation of propionyl-CoA was largely unaffected, indicating a preference for propionyl-CoA under these conditions (Fig. 1d).

### LCC adopts a modular and highly dynamic structural architecture

To capture distinct states along the LCC catalytic cycle, we determined cryo-EM structures of cytosolic LCC in the presence of C16-CoA alone, a mixture of C3- and C16-CoA, or C16-CoA with ATP to induce turnover. A composite cryo-EM reconstruction of C16-CoA-bound LCC at an average resolution of 3.1 Å revealed a modular architecture comprising a central carboxyltransferase (CT) core tethered to two mobile biotin carboxylase (BC) modules via the flexible, non-catalytic linker subunit AccE5, representing a pre-activation state in the absence of ATP (Fig. 2a, Supplementary Fig. 3-4, Supplementary Table 1-3). This reconstruction defines an overall stoichiometry of eight AccA3 subunits, two AccD4 subunits, four AccD5 subunits, and two AccE5 subunits.

**Figure 2.**
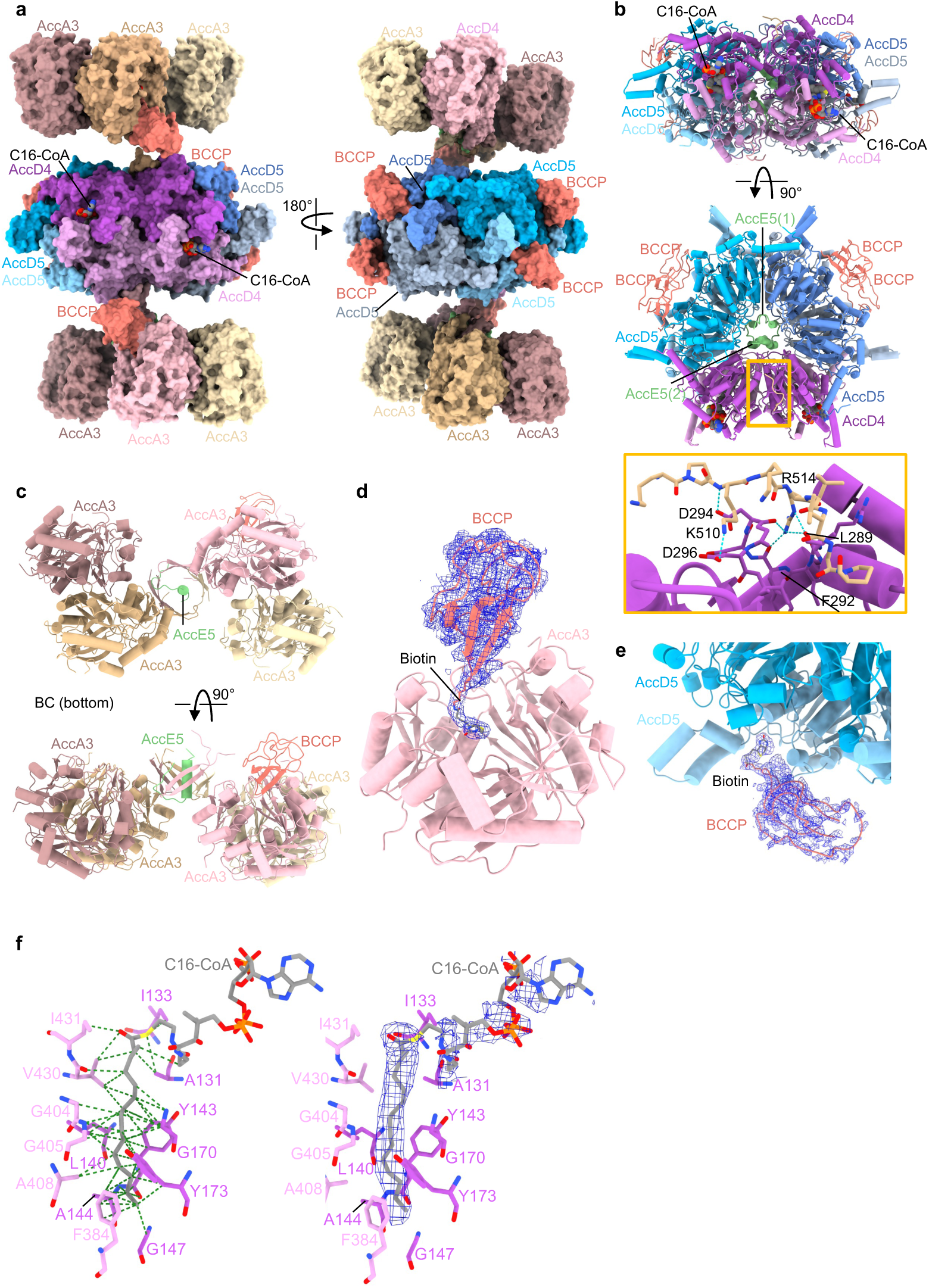
Architecture of the LCC complex in pre-activation state bound to palmitoyl-CoA. **a,** Atomic model of the LCC complex (surface representation) build into the composite cryo-EM map (supplementary Fig. 4). Central carboxyltransferase (CT) core is flanked by two biotin carboxylase (BC) modules (top and bottom) connected by flexible AccE5 linkers (not visible). Palmitoyl-CoA is bound in the AccD4 (purple) active site. Two BCCPs (salmon) engage the BC modules (left). A view rotated by 180° shows four BCCPs positioned near the AccD5 subunit of the CT core (right). **b,** Side and top views of the atomic model for the CT module. The C-terminal helix-turn-helix of AccE5 (green) anchors the CT core. An AccA3 loop (brown) interacts with AccD4 (purple). **c,** Top and side views of the atomic model for the BC module. The N-terminal α-helix of AccE5 (green) anchors the AccA3 tetramer. **d,** Pre-activation BCCP (blue mesh density) with biotinylated Lys564 (blue mesh density) positioned in the ATP-binding pocket of AccA3 (salmon). **e,** CT-proximal BCCP (blue mesh density) transiently samples the AccD5 (blue) substrate entry. **f,** Representative close-up view of the AccD4 active site engaging C16-CoA predominantly through van der Waals interactions (green dashes) with highlighted residues. The right panel shows cryo-EM density for C16-CoA (blue mesh).

### The CT module harbours six active sites

The CT module, which catalyses transfer of CO₂ from carboxybiotin to acyl-CoA substrates, forms a pseudo-D3-symmetric double-ring assembly composed of one AccD4 dimer and two AccD5 dimers arranged as a trimer of dimers (Fig. 2b). AccD4 and AccD5 both adopt the canonical crotonase fold and are highly similar in structure (Z-score 22.8; RMSD 1.11 Å over 494 residues; Supplementary Fig. 5), giving rise to this pseudo-symmetry. This architecture generates six carboxyltransferase active sites, with two located at the AccD4-AccD4 homodimer interfaces and four at the AccD5-AccD5 homodimer interfaces, where N- and C-terminal domains from adjacent protomers form the catalytic centers.

### AccE5 tethers the BC and CT modules through a bipartite anchoring mechanism

AccE5 links the CT core to the BC modules through two interfaces. Its C-terminal helix-turn-helix motif inserts into the CT-ring lumen and contacts AccD4 and AccD5 protomers (Fig. 2b). At its N terminus, AccE5 anchors to each BC tetramer through a helix-in-a-β-cage interface, in which two β-strands from each AccA3 protomer and one from AccE5 form a nine-stranded β-barrel enclosing the AccE5 helix (Fig. 2c). A long intrinsically disordered region, unresolved in the density, separates the two interfaces and permits flexible BC-CT pivoting required for BCCP shuttling.

### BCCP occupies distinct BC-anchored and CT-proximal states before activation

Within each BC module, AccA3 assembles as a tetramer (Fig. 2c) adopting the conserved ATP-grasp fold. Each protomer comprises a catalytic BC core, a flexible ATP cap domain (residues 143-211 unresolved), and a biotin carboxyl carrier protein (BCCP) domain that carries the biotin prosthetic group and is tethered by a flexible, mostly unresolved linker that enables shuttling of the BCCP domain between the BC module and the CT core.

One BCCP is resolved per BC tetramer, with the biotinylated Lys564 inserted into the ATP-binding pocket in a pre-activation pose poised for ATP-dependent carboxylation (Fig. 2d). A short linker segment contacts the AccD4 dimer, positioning this BC-anchored BCCP proximal to the CT module, primarily via residues K512 and R517, which form salt bridges and hydrogen bonds with AccD4 (Fig. 2b).

In addition to these BC-anchored BCCPs, four further BCCPs are observed near the AccD5-facing surface of the CT core (Fig. 2a). These CT-proximal BCCPs show partial density, whereas the uncarboxylated biotin moiety is well defined as it samples the active-site entrance, consistent with transient, non-productive docking (Fig. 2e). Linker density is unresolved for these CT-bound BCCPs, precluding assignment to specific AccA3 subunits, and two additional BCCPs remain undetected, consistent with high mobility. Notably, despite the presence of C16-CoA in AccD4, no BCCP engages the AccD4 active site under ATP-free conditions, indicating preferential sampling between the BC module and AccD5 prior to activation.

### Structural basis of C16-CoA recognition by AccD4

The CT-core, resolved at 2.3 Å resolution (Supplementary Fig. 3), shows that the acyl-chain tunnel accommodates a well-defined C1-C16 segment of the substrate C16-CoA (Fig. 2f; Supplementary Fig. 6), whereas the CoA moiety is poorly resolved, with weak density observed for the S1P-C7P segment and fragmented density for the adenine ring, indicating flexibility of the CoA moiety. AccD4 engages C16-CoA primarily through a tight network of van der Waals interactions surrounding the acyl chain.

### ATP turnover drives BCCP redistribution and BC-CT rearrangements

To examine LCC under ATP-turnover conditions, we analyzed C16-CoA-bound LCC in the presence of ATP. A composite reconstruction refined to an average resolution of 2.9 Å reveals pronounced ATP-dependent conformational rearrangements of the complex (Fig. 3a-c; Supplementary Fig. 7-8; Supplementary Tables 4-6). Comparison with the ATP-free pre- activation state shows that ATP binding is accompanied by rotations of the two BC tetramers by ∼65° and ∼62°, respectively, relative to the CT core (Fig. 4).

**Figure 3.**
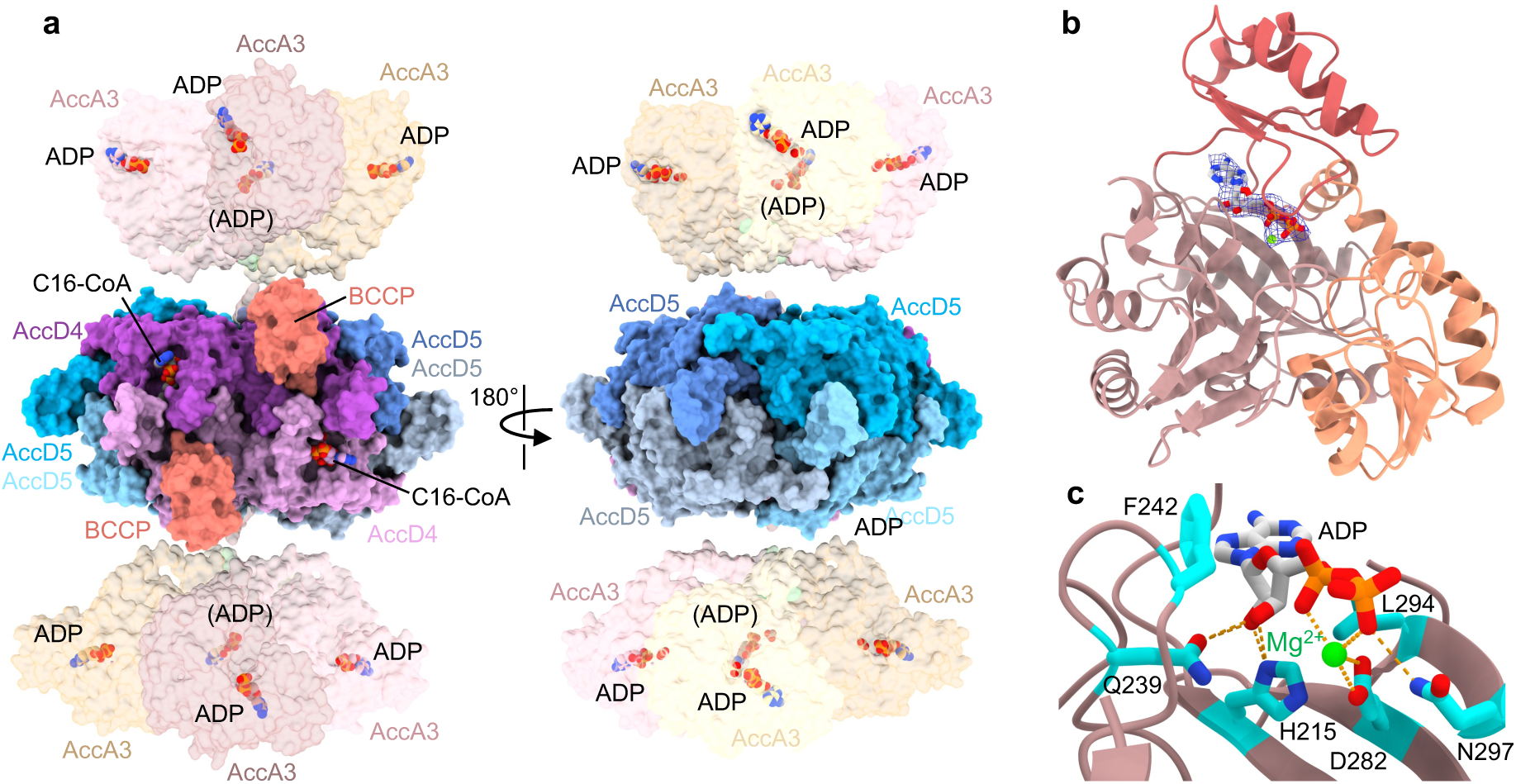
Architecture of the LCC complex under ATP turnover conditions. **a,** Atomic model of the LCC complex (surface representation) built into the composite cryo-EM map (Supplementary Fig. 8), with C16-CoA bound in the AccD4 active sites (purple), two BCCP domains (salmon) positioned proximal to AccD4, and ADP-Mg²⁺ bound to AccA3. A view rotated by 180° is shown on the right. **b,** Atomic model of an AccA3 monomer bound to ADP (blue mesh density) and Mg²⁺ (green sphere). **c,** Interactions between ADP, Mg²⁺, and active-site residues.

**Figure 4.**
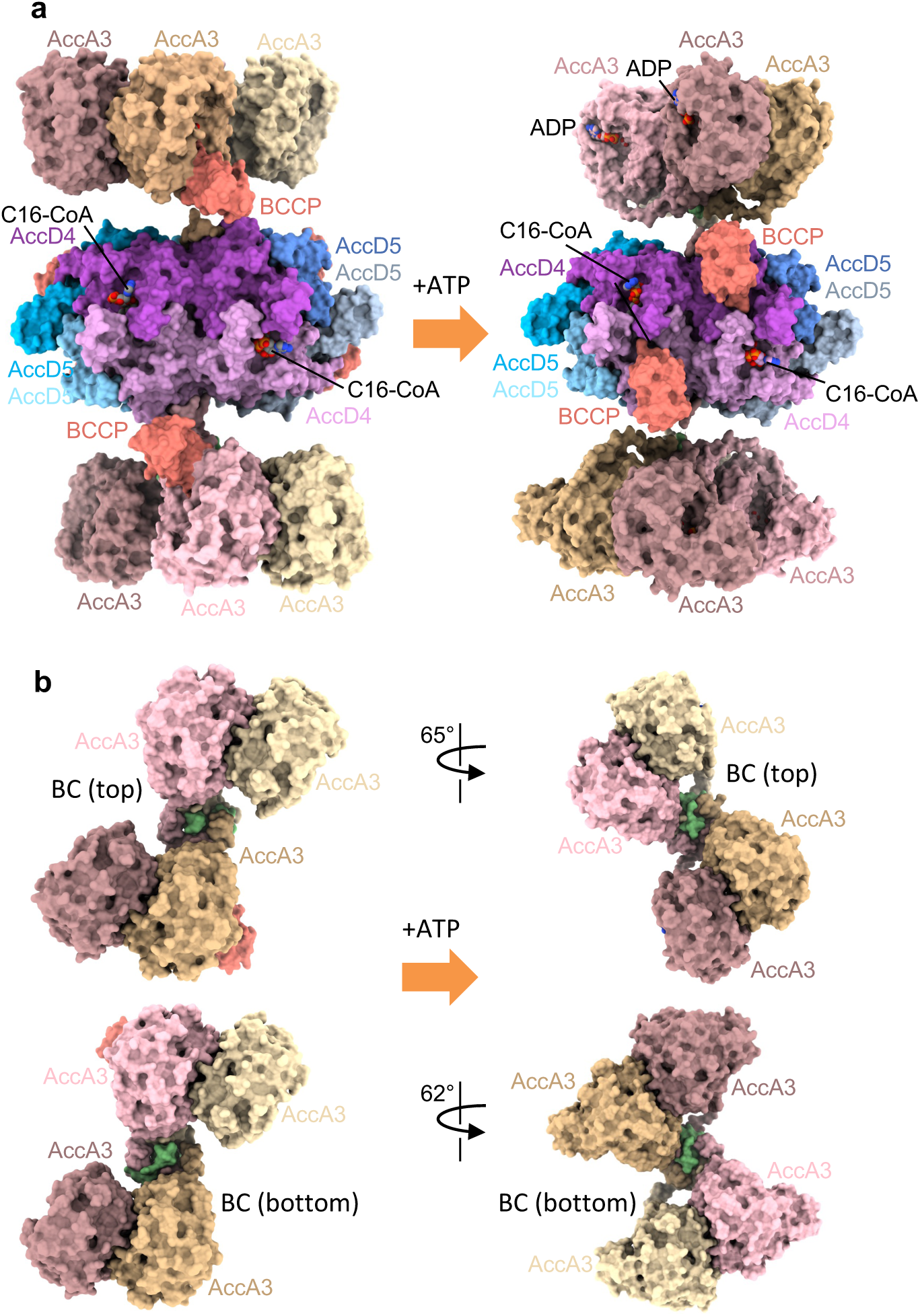
ATP-dependent rotation of BC modules relative to the CT core. **a,** Surface representations of the LCC atomic models in the pre-activation state, with C16-CoA bound to AccD4 in the absence of ATP (left), and under ATP turnover conditions (right), in which BCCP (salmon) associates with the C16-CoA-bound AccD4 site. **b,** Comparison of the two states reveals ATP-dependent rotations of ∼65° and ∼62° for the top and bottom BC modules, respectively, relative to the CT core.

Concomitant with this BC-CT rearrangement, BCCPs dissociate from the BC tetramers, where density consistent with bound ADP is observed in the AccA3 active sites, indicating a post- carboxylation state of the BC module (Fig. 3b-c). This density was assigned as ADP coordinated by Mg²⁺ based on structural superposition with the biotin carboxylase from *Escherichia coli* resolved in complex with ADP-Mg²⁺ (PDB ID 3RV3; RMSD 0.92 Å over 344 residues; Supplementary Fig. 9). Under turnover conditions, BCCP domains instead associate with the C16-CoA-engaged AccD4 sites (Fig. 3a), whereas no BCCP density is detected at AccD5, indicating preferential association with substrate-engaged AccD4 under ATP turnover conditions.

Compared with the ATP-free state, BCCP density at AccD4 is markedly more diffuse, consistent with rapid shuttling between the BC and CT modules. The biotinylated linker lysine is not resolved, suggesting fast insertion and withdrawal cycles during catalysis. Consistent with this dynamic behavior, no density corresponding to a carboxylated C2 position of C16-CoA is observed (Supplementary Fig. 10). Together, these observations link ATP-dependent BC rotation to large-scale reorganization of BCCP positioning during the LCC catalytic cycle.

### Long- and short-chain substrates are simultaneously accommodated by AccD4 and AccD5

Next, we analyzed the ATP-free pre-activation state of LCC in the presence of both C3- and C16-CoA (Supplementary Fig. 11). The resulting reconstructions reveal simultaneous binding of C16-CoA at AccD4 and C3-CoA at AccD5, yielding an overall architecture that closely resembles the ATP-free, C16-CoA-bound pre-activation state. For the majority of particles, only the CT core could be resolved to high resolution (2.2 Å), whereas the flexible BC tetramers remained poorly defined. However, a minor particle population yielded a composite reconstruction (Supplementary Fig. 12; Supplementary Tables 7-9). In this structure, four BCCP domains localize near the four C3-CoA-engaged AccD5 active sites (Fig. 5a), while no BCCP density is detected at the two C16-CoA-bound AccD4 sites, mirroring the C16-CoA-only pre-activation state of LCC. Both BC tetramers tilt toward the CT core and adopt a more pronounced inclination than observed in the C16-CoA-only pre-activation state. Within each BC tetramer, one BCCP and its associated linker contact an AccD4 protomer. The biotinylated Lys564, however, remains oriented toward the BC active site and poised for carboxylation (Supplementary Fig. 13), indicating that BCCP engagement with AccD4 does not occur prior to ATP-dependent activation and is consistent with the pre-activation configuration observed for C16-CoA-bound LCC (Fig. 2d).

**Figure 5.**
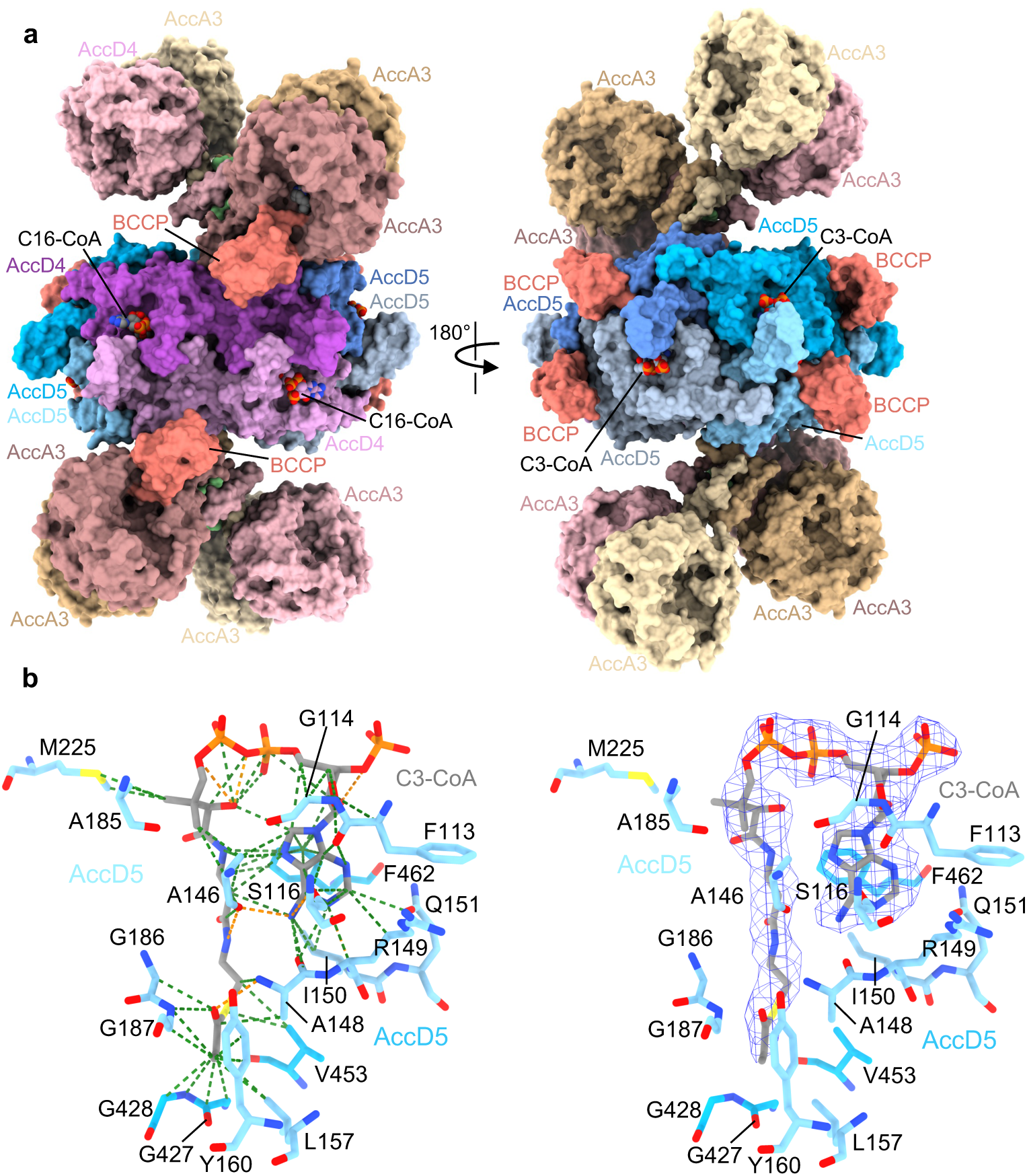
Architecture of LCC and structural basis of C3-CoA and C16-CoA binding. **a,** Atomic model of the LCC complex (surface representation) built into the composite cryo-EM map (Supplementary Fig. 11), with C3-CoA bound in the AccD5 active site (blue shades) and C16-CoA bound in the AccD4 active site (purple shades). Two BCCP domains (orange) engage the biotin carboxylase (BC) modules (left), whereas four BCCPs localize near the AccD5-facing surface of the carboxyltransferase (CT) core (right). **b,** Detailed view of C3-CoA interactions within the AccD5 active site. Hydrogen bonds are shown as orange dashed lines and van der Waals interactions as green dashed lines. The right panel shows cryo-EM density for C3-CoA (blue mesh).

### Structural basis of C3-CoA recognition by AccD5

We next examined how the CT module engages the short-chain substrate C3-CoA at AccD5. Density for C3-CoA is well resolved, with the CoA moiety adopting a compact, bent conformation (Fig. 5b). The adenine base is coordinated by hydrogen bonds to backbone carbonyls of residues S116, A146, A148, and I150, and is further positioned by an aromatic interaction with F462. The C1 carbonyl of the acyl chain is stabilized by hydrogen bonds from A148 and G187.

### Structural determinants of substrate selectivity in AccD4 and AccD5

To understand how the two CT subunits discriminate between acyl-CoA substrates, we compared the active-site architectures of C16-CoA-bound AccD4 and C3-CoA–bound AccD5. Superposition of the two dimers shows that placement of a C16 acyl chain into the AccD5 active site would result in steric overlap with surrounding residues, indicating that the AccD5 pocket cannot accommodate long-chain substrates (Fig. 6a).

**Figure 6.**
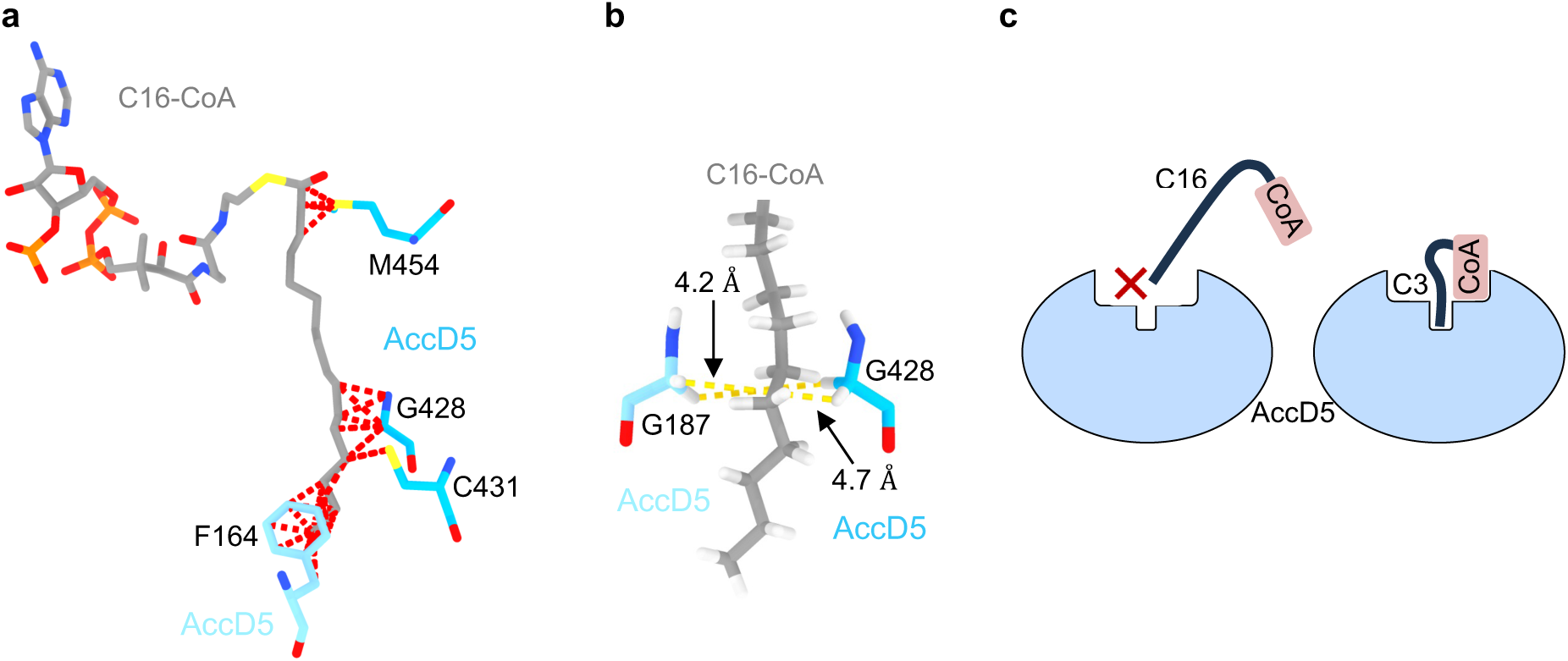
Structural basis of substrate selectivity in AccD4 and AccD5. **a,** Superposition of the C16-CoA-bound AccD4 dimer with the C3-CoA-bound AccD5 dimer. The position of the C16 acyl chain relative to the AccD5 active site is shown, with steric overlap between the acyl chain and surrounding AccD5 residues indicated by red dashed lines (AccD5 dimer shown in blue). The CoA moiety and part of the acyl chain are omitted for clarity. **b,** Close-up view of the AccD5 active site formed at the dimer interface, highlighting residues G187 (monomer 1) and G428 (monomer 2), which define a constriction within the acyl-chain tunnel (AccD5 shown in blue). The CoA moiety and part of the acyl chain are omitted for clarity. **c,** Schematic summary of substrate selectivity illustrating that AccD5 (blue) accommodates C3-CoA but not C16-CoA.

Inspection of the AccD5 active site reveals the structural basis for this exclusion. The acyl-chain tunnel contains a constriction of 4.2-4.7 Å at the interface between opposing protomers, defined by residues G187ʹ and G428 (Fig. 6b). This opening lies below the ∼5.2 Å steric minimum required to accommodate an acyl chain of ∼3 Å diameter, thereby precluding binding of long-chain acyl-CoA substrates.

AccD4 engages C16-CoA predominantly through extensive van der Waals interactions along the acyl chain, with comparatively few contacts to the CoA moiety (Fig. 2f; Supplementary Fig. 6). By contrast, AccD5, whose binding pocket lacks sufficient volume for long-chain substrates, binds C3-CoA through limited interactions with the short acyl chain but primarily through contacts with the CoA group (Fig. 5b). These differences in pocket accessibility and substrate engagement explain why AccD4, but not AccD5, can accommodate long-chain acyl-CoA substrates (Fig. 6c).

### Conserved catalytic architecture with differential substrate positioning

Structural comparison of AccD4 and AccD5 with PccB from *Streptomyces coelicolor* bound to C3-CoA and biotin (PDB ID 1XNY) identifies two conserved glycine-based oxyanion holes in both AccD4 and AccD5 that are required for carboxyl transfer (Fig. 7a-b). Oxyanion hole 1 (G169ʹ-A170ʹ in AccD4; G186ʹ-G187ʹ in AccD5) is positioned to stabilize the substrate enolate following α-proton abstraction, whereas oxyanion hole 2 (G405-G406 in AccD4; G428-A429 in AccD5) is positioned to stabilize the carboxylate of carboxybiotin during CO₂ transfer. In C3- CoA-engaged AccD5, the C2 (α-carbon) of the bound substrate- the site of carboxylation-aligns with these conserved oxyanion holes (Fig. 7c). Despite this conserved catalytic architecture, the α-carbon of C16-CoA in AccD4 remains more than 10 Å from the catalytic center (Fig. 7d), compared with the ∼3 Å separation observed in transfer-competent PccB structures. Thus, long-chain carboxylation by AccD4 requires an additional conformational rearrangement to bring the acyl substrate into a transfer-competent geometry, which is not resolved in the present structures.

**Figure 7.**
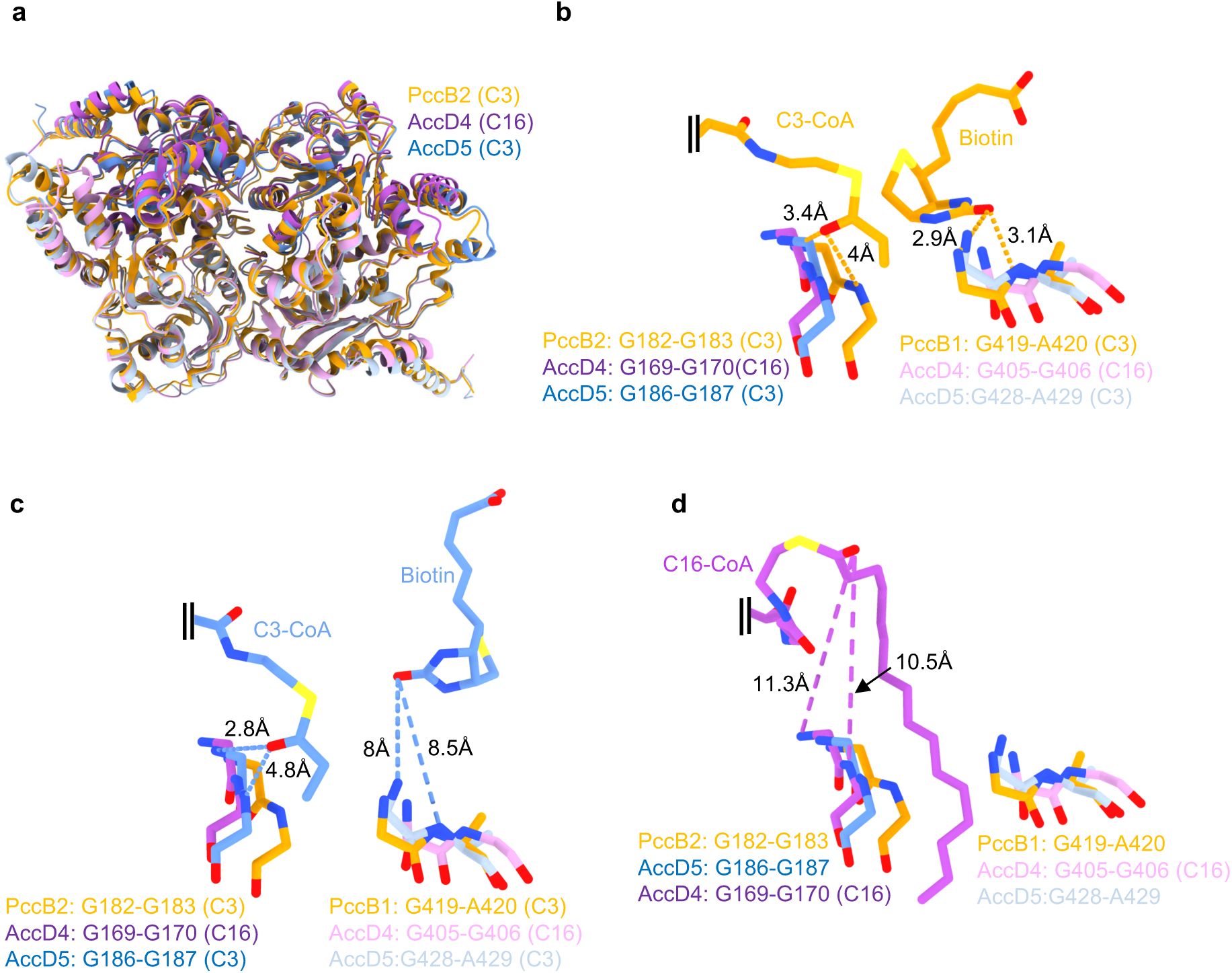
Comparison of ligand positioning relative to conserved oxyanion holes in PccB, AccD4, and AccD5. **a,** Structural superposition of the C3-CoA-bound PccB dimer from *Streptomyces coelicolor* (PDB 1XNY) with the C3-CoA-bound AccD5 dimer (blue) and the C16-CoA-bound AccD4 dimer (purple). **b-d,** Close-up views of the active sites after structural superposition, showing conserved oxyanion-hole residues (stick representation) in equivalent positions. **b,** PccB with C3-CoA and biotin (orange); **c,** AccD5 with C3-CoA and biotin (blue); **d,** AccD4 with C16-CoA (purple). Distances between ligands and oxyanion-hole residues are indicated.

### Structural organization of an AccD5-AccE5 core and its association with AccA3 modules

In addition to LCC, particle classification revealed a distinct acyl-CoA carboxylase assembly centered on AccD5 and the ε-subunit AccE5. In datasets collected without added ATP, this complex resolves as a hexameric AccD5 carboxyltransferase core lacking associated biotin carboxylase (BC) modules (Fig. 8a-b; Supplementary Fig. 14b; Tables 2-3). In the presence of C3-CoA, density for bound substrate is observed at the AccD5 active sites, with the CoA moiety adopting a compact conformation comparable to that observed for AccD5 within LCC (Supplementary Fig. 15b, 18; Tables 8-9), indicating that the AccD5-centered core can directly engage short-chain substrates independently of associated BC modules. An apo form of the same assembly is observed in parallel, lacking substrate density (Fig. 8a). Although density is present at the expected AccE5 position in the hexameric AccD5 assembly, it is fragmented and precludes confident modeling.

**Figure 8.**
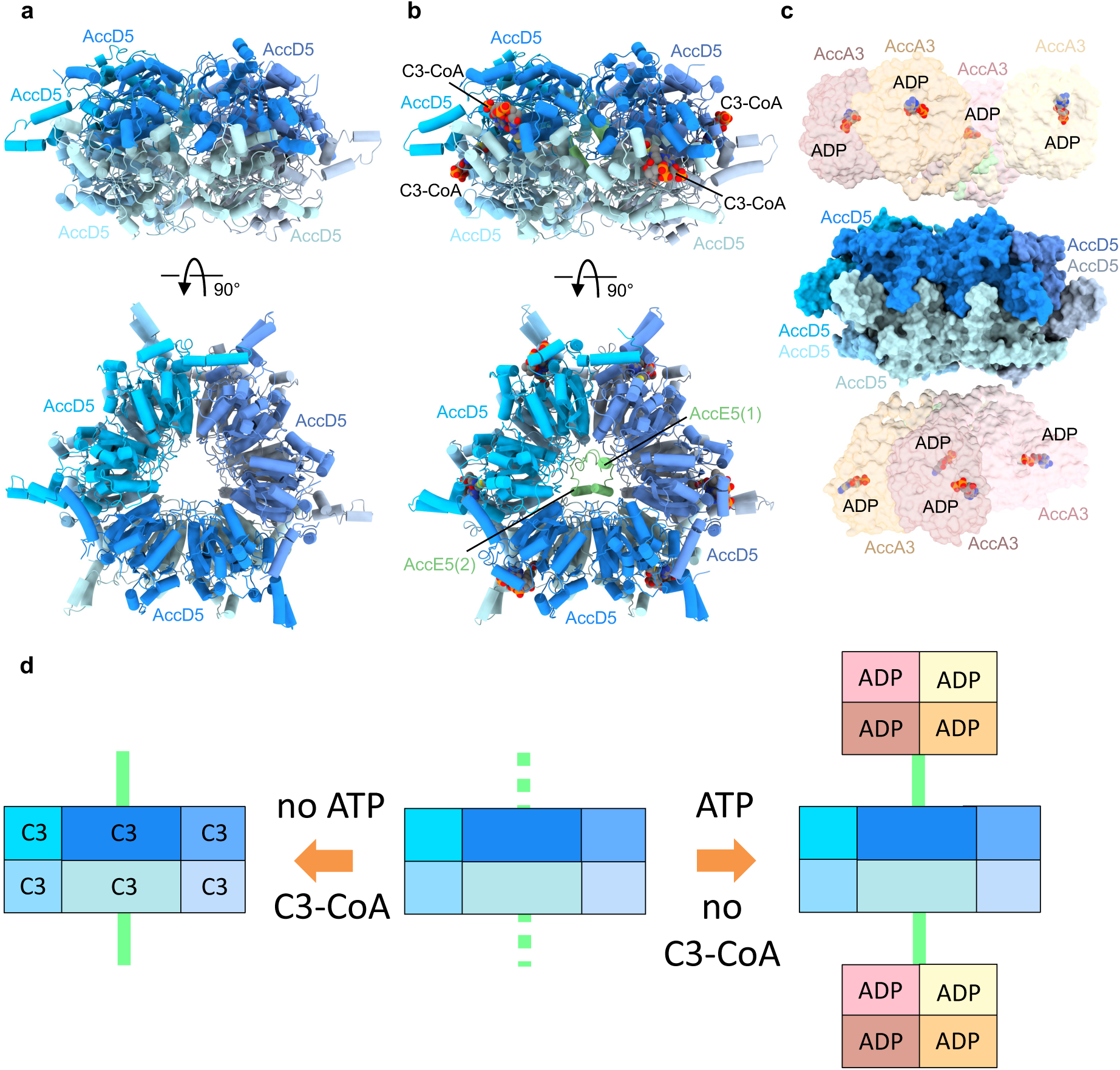
Cryo-EM reconstructions of an AccD5-centered carboxyltransferase assembly. **a,** Top and side views of the apo AccD5-centered assembly resolved in datasets collected without ATP and C3-CoA. Density at the expected AccE5 position is fragmented and was not modelled. **b,** Top and side views of the AccD5-centered assembly with C3-CoA bound at the active sites in datasets collected without ATP. **c,** Under ATP-turnover conditions, an expanded assembly is observed in which AccA3-containing biotin carboxylase (BC) modules associate with the AccD5–AccE5 core, yielding an 8AccA3:6AccD5:2AccE5 complex corresponding to the short-chain acyl-CoA carboxylase activity previously referred to as ACCase 5. **d,** Schematic representation of the AccD5-centered assemblies identified in this study under the indicated conditions. The carboxyltransferase core forms a homohexameric AccD5 scaffold that associates with biotin carboxylase (BC) modules, illustrating the modular organization, also observed in LCC.

Under ATP-turnover conditions, a related but expanded assembly is observed in which AccA3-containing BC modules associate with the AccD5-AccE5 core, yielding an 8AccA3:6AccD5:2AccE5 complex (Supplementary Fig. 16-17; Tables 5-6). This assembly structurally defines the short-chain acyl-CoA carboxylase activity previously referred to as ACCase 5 in the literature^17–19^ . A composite reconstruction refined to ∼3.2 Å resolution reveals AccA3 arranged in tetrameric BC modules analogous to those in LCC, with density for bound ADP present in the AccA3 active sites (Fig. 8c; Supplementary Fig. 19). In this state, no substrate or BCCP density is detected, consistent with a BC-engaged post-reaction configuration. To our knowledge, this represents the first structural visualization of ACCase 5 within an intact assembly.

Together, these observations provide direct structural evidence for a combinatorial assembly mechanism of mycobacterial acyl-CoA carboxylases. Distinct carboxyltransferase cores-either the AccD4/AccD5 heterohexamer of LCC or the AccD5 homohexamer corresponding to ACCase 5-can associate with biotin carboxylase modules to generate alternative holoenzyme architectures. This modular organization is consistent with prior biochemical reconstitution and interaction studies and establishes structural principles governing assembly within the mycobacterial acyl-CoA carboxylase network.

## Discussion

In this work, we define the architecture of the mycobacterial long-chain acyl-CoA carboxylase and reveal the structural basis for dual substrate selectivity. The LCC architecture differs in both stoichiometry and inter-module connectivity from that of known bacterial and eukaryotic acyl-CoA carboxylase complexes^16,22–24^. Whereas canonical acetyl-CoA carboxylases adopt a symmetric 6:6 organization^22–25^ in which six BC domains pair with six CT active sites, LCC is asymmetric, comprising eight BC active sites arranged as two AccA3 tetramers but only six CT sites formed by a 2:4 AccD4/AccD5 heterohexamer. This 8:6 mismatch is unprecedented among biotin-dependent carboxylases. A second distinguishing feature of LCC is the presence of two AccE5 subunits that flexibly tether each BC module to the CT ring. No comparable anchoring mechanism has been described for other carboxylases, in which BC and CT assemblies associate through direct, symmetric interfaces. This asymmetric architecture is consistent with the expanded substrate range of LCC, which accommodates both short-chain and very long-chain acyl-CoA substrates via the specialized AccD5 and AccD4 active sites, respectively. In contrast, symmetric ACCs typically carboxylate a single substrate class, establishing LCC as a structurally distinct member of the ACC family. Despite its unconventional architecture, LCC undergoes conformational transitions that follow principles established for bacterial biotin-dependent carboxylases. ATP binding promotes redistribution of BCCP from the ADP bound BC module toward the CT region, consistent with long-range biotin shuttling described for pyruvate carboxylase^26^.

Beyond LCC, our structures reveal that AccD5 forms a stable homohexameric carboxyltransferase core that associates with AccE5 and, under turnover conditions, engages AccA3-containing BC modules to generate an alternative 8AccA3:6AccD5:2AccE5 assembly. This assembly structurally defines the short-chain acyl-CoA carboxylase activity previously referred to as ACCase 5^17–19^. While AccD5-dependent short-chain activities had been inferred from biochemical reconstitution and interaction studies ^10,17–19^, their structural organization was unknown. The present structures provide this missing architectural framework.

The coexistence of an AccD4/AccD5 heterohexameric CT core in LCC and an AccD5 homohexameric core corresponding to ACCase 5 demonstrates that distinct carboxyltransferase cores can associate with shared biotin carboxylase modules to generate alternative holoenzyme architectures. These findings provide direct structural evidence for a combinatorial assembly mechanism of mycobacterial acyl-CoA carboxylases, in which CT core identity dictates substrate range and functional specialization. Rather than existing as fixed holoenzymes, mycobacterial carboxylases function as modular assemblies capable of reorganizing around alternative CT scaffolds.

We further show that the short-chain substrates C2-CoA and C3-CoA compete for AccD5-mediated carboxyltransferase sites, with propionyl-CoA preferentially retained over acetyl-CoA. These data indicate that short-chain substrate utilization by mycobacterial acyl-CoA carboxylase assemblies depends on relative intracellular availability and access to AccD5-containing CT cores.

Despite differences in substrate range, the catalytic architecture of the CT module is conserved. Both AccD4 and AccD5 harbor two glycine-based oxyanion holes characteristic of biotin-dependent carboxylases. In the C3-CoA-bound AccD5 structure, the distance between the C2 carbon of the substrate and oxyanion hole 1 is compatible with CO₂ transfer from carboxybiotin, comparable to transfer-competent PccB structures. In contrast, in all captured states the α-carbon of C16-CoA in AccD4 remains more than 10 Å from the catalytic center, indicating that these conformations are not transfer-competent and that long-chain carboxylation requires a short-lived rearrangement not resolved here.

Although we did not visualize the transient carboxylated biotin intermediate under turnover conditions, we observed sampling of the CT active site by uncarboxylated biotin in pre-activation states. In these configurations, the separation between biotin and oxyanion hole 2 exceeds that required for productive transfer^27^, consistent with previous studies showing that only proximal biotin conformations support carboxyl transfer^28^ ^29^.

Finally, the core architecture of LCC is conserved across mycolic-acid-producing Corynebacteriales. AlphaFold-based modeling supports conservation of both BC and CT modules in major pathogens, including *M. tuberculosis*, *M. abscessus*, and *M. leprae*, as well as in the industrial species *Corynebacterium glutamicum*, indicating that the mechanistic principles defined here are broadly applicable.

Together, our findings establish LCC and ACCase 5 assemblies as modular components of a combinatorial acyl-CoA carboxylase platform and define the structural basis of substrate selectivity and assembly-specific function in mycobacteria.

## Methods

### Bacterial strains and growth conditions

*Mycobacterium smegmatis* mc²155 was transformed with the plasmid pMyNT empty vector under the control of the acetamidase promoter. Cultures were grown in Luria-Miller (LB) broth supplemented with 0.2% (v/v) glycerol at 180 rpm. Expression was induced with 0.2% (w/v) acetamide at an optical density at 600 nm (OD₆₀₀) of 0.5, and cells were harvested at a final OD₆₀₀ of 2.0-2.1. Cells were collected by centrifugation and washed three times with phosphate-buffered saline (PBS). As a control, uninduced pMyNT empty-vector cultures were processed in parallel under identical conditions.

### Purification of the LCC complex

Cell pellets from induced and uninduced pMyNT empty-vector cultures were resuspended at a 1:5 (w/v) ratio in buffer A (300 mM NaCl, 50 mM HEPES, 1 mM DTT, pH 8.0) supplemented with EDTA-free protease inhibitors (Roche) and lysozyme (Roche). Cells were lysed by three passages through an Emulsiflex-C3 homogenizer (Avestin), and unlysed material was removed by centrifugation at 10,000 × g for 15 min at 4 °C.

The soluble fraction was separated by ultracentrifugation at 45,000 rpm for 1 h at 4 °C using a 45Ti rotor and applied directly to a pre-equilibrated Strep-Tactin™ XT 4Flow™ column (IBA). The column was washed with buffer A until a stable UV baseline was achieved, and bound proteins were eluted with buffer B (buffer A supplemented with 50 mM biotin, pH 8.0). Eluted samples were further purified by size-exclusion chromatography on a Superose 6 Increase 10/300 GL column (GE Healthcare) equilibrated in buffer C (300 mM NaCl, 20 mM HEPES, 1 mM DTT, pH 8.0). Fractions were analyzed by SDS-PAGE and BN-PAGE followed by colloidal Coomassie Blue staining.

### In-gel digestion and nano-LC-MS workflow

Proteins were separated by reducing SDS-PAGE, stained with Coomassie Blue, and excised bands were cut into ∼1 mm³ pieces. In-gel digestion was performed as described by Li *et al*^30^. Gel pieces were washed repeatedly with acetonitrile (ACN) and water to remove SDS, reduced with 2 mM DTT (30 min, 37 °C), alkylated with 5 mM iodoacetamide (30 min, room temperature, dark), and digested overnight at 37 °C with sequencing-grade trypsin (10 ng/µl; Promega) in 50 mM ammonium bicarbonate containing 1 mM CaCl₂. Peptides were extracted with ACN and ACN/ammonium bicarbonate, dried by vacuum centrifugation, resuspended in 97:3:0.1 (H₂O:ACN:formic acid), and desalted using C18 StageTips following standard protocols.

Desalted peptides were analyzed by nanoLC-MS/MS on an UltiMate 3000 RSLCnano system coupled to a Q Exactive HF mass spectrometer (Thermo Fisher Scientific). Peptides were separated using a trap-elute configuration with a nanoEase M/Z Symmetry C18 trap column (180 µm × 20 mm) and a nanoEase M/Z HSS C18 T3 analytical column (75 µm × 250 mm), maintained at 40 °C. Peptides were eluted over an 85 min gradient from 1% to 40% mobile phase B (0.1% formic acid in acetonitrile) at 0.3 µl/min.

The mass spectrometer was operated in positive-ion, data-dependent acquisition mode. Full MS scans were acquired at 60,000 resolution (m/z 200), followed by up to 10 MS/MS scans at 15,000 resolution using higher-energy collisional dissociation (normalized collision energy 28). Dynamic exclusion was set to 20 s, and singly charged and unassigned ions were excluded. Instrument performance was monitored using routine quality control injections.

Raw data were processed using MaxQuant (v1.6.17.0) with label-free quantification and match-between-runs enabled. Searches were performed against the UniProt reviewed database supplemented with common contaminants.

### Cryo-EM sample preparation and data collection

To generate palmitoyl-CoA-bound LCC, LCC was incubated with 500 μM palmitoyl-CoA for 2h before vitrification. And for the palmitoyl-CoA bound complex under ATP turnover condition, samples were incubated with 500 μM palmitoyl-CoA, 25mM NaHCO_3,_ 10mM ATP, and 10mM MgCl_2_ for 2h before vitrification. Propionyl- and palmitoyl-CoA-bound LCC was generated by addition of 1 mM propionyl-CoA and 500 μM palmitoyl-CoA.

Cryo-EM grids were prepared by applying 3.5 µl of sample to glow-discharged gold grids (R1.2/1.3, 300 mesh, Ultrafoil). Grids were blotted for 3.5 s with a blot force of 10 using a Vitrobot Mark IV (Thermo Fisher Scientific) operating at 4 °C and 100% relative humidity, followed by rapid vitrification in liquid ethane. Data were acquired on a Titan Krios G1 transmission electron microscope (Thermo Fisher Scientific) operating at 300 kV, equipped with a Gatan K3 direct electron detector and a BioQuantum energy filter (Gatan) with a 20eV slit width. For the palmitoyl-CoA, palmitoyl-CoA under ATP turnover condition, and the mixture of propionyl-CoA and palmitoyl-CoA, a total of 14,716, 15,130, and 14,974 movies, respectively, were collected in electron counting mode using aberration-free image shift as implemented in EPU (Thermo Fisher Scientific). Movies were recorded at a nominal magnification of 105,000×, corresponding to a calibrated pixel size of 0.836 Å. Each movie was acquired over 60 frames with a total dose of 60 e⁻/Å² and a defocus range of −0.8 to −2.0 µm.

### Cryo-EM data processing

All the cryo-EM data were processed using CryoSPARC v4.5.3 and v4.7.1 (Structura Biotechnology Inc.). Patch-based motion correction and contrast transfer function (CTF) estimation were performed on all acquired micrographs^31^. Micrographs were manually curated, and a subset was used for initial particle picking. Manually selected particles were subjected to 2D classification, and high-quality class averages were used as templates for automated particle picking.

For the LCC complex incubated with palmitoyl-CoA cryo-EM data (Supplementary Fig.3), particles were extracted with a box size of 512 pixels and Fourier cropped to 220 pixels. Multiple rounds of 2D classification were performed to remove junk particles. An initial ab initio reconstruction was generated from selected particles, followed by iterative heterogeneous refinement to resolve compositional heterogeneity. Particles corresponding to the best-resolved classes were re-extracted with a box size of 480 pixels, Fourier cropped to 416 pixels, and subjected to non-uniform refinement^32^. This yielded consensus map at resolution 2.15 Å measured by gold-standard Fourier shell correlation at 0.143 criterion. The resulting map exhibited reduced resolution at the distal ends of the complex (AccA3 at top and bottom), attributed to conformational flexibility. To further improve the reconstruction, the non-uniform refinement map was used for mask generation. Density subtractions were employed to isolate the AccA3 at top, 4AccD5:2AccD4:2AccE5 at centre, and AccA3 at bottom regions. Focused refinement of the centre region (4AccD5:2AccD4:2AccE5) yielded a locally refined map at 2.16 Å resolution.

Given the substantial conformational variability, 3D variability analysis^33^ was performed on the 4AccD5:2AccD4:2AccE5, and the resulting principal components were analyzed in ChimeraX v1.9. Based on the principal component analysis, particles were subjected to 3D variability display (cluster mode) job in cryoSPARC v4.5.3 (Structura Biotechnology Inc.), and the best-resolved classes were subjected to non-uniform refinement. This particle sorting strategy resulted in two distinct complexes 4AccD5:2AccD4:2AccE5 and 6AccD5:2AccE5. The final reconstruction result in resolution 2.3 Å for 4AccD5:2AccD4:2AccE5 and 2.9 Å for 6AccD5:2AccE5 (supplementary Fig. 14a, b), respectively, which were subsequently used for atomic model building. To resolve flexible AccA3, direct local refinement from particle subtractions of the flexible AccA3 at top and bottom regions did not produce substantial improvements. To solve this problem, the subtracted particles corresponding to these regions were subjected to ab initio reconstruction to generate three initial models. These models were used in heterogeneous refinement to classify the particle population. The best-resolved classes were further subjected to non-uniform refinement, resulting in improved density maps for the top and bottom regions. Final resolutions, determined at the gold-standard Fourier shell correlation (FSC) 0.143 criterion, were 3 Å and 3 Å, respectively (Supplementary Fig.14c,d), and were subsequently used to guide atomic model building. Further, to get full map of both the complexes, 3D classification was performed on full set of particles from non-uniform refinement job. Class with 65,445 particles was selected where full LCC complex (4AccD5:2AccD4:8AccA3:2AccE5) could be resolved. These particles subjected to non-uniform refinement and further local refinement to get high resolution of AccA3 at top, 4AccD5:2AccD4:2AccE5 at centre, and AccA3 at bottom. The resulting locally refined maps were combined to generate a composite reconstruction of the full LCC complex (Supplementary Fig.4). We could not resolve the full reconstruction of other complex.

For the LCC complex incubated with palmitoyl-CoA, ATP, MgCl2, and NaHCO3 cryo-EM data (Supplementary Fig.7,16) particles were extracted with a box size of 480 pixels, Fourier cropped to 220 pixels. Multiple rounds of 2D classification were performed to remove junk particles. An initial ab initio reconstruction was generated from selected particles, followed by iterative heterogeneous refinement to resolve compositional heterogeneity. Particles of overexpressed views were removed by using the ’rebalance orientation’ job. Balanced particles were re-extracted with a box size of 480 pixels, Fourier cropped to 416 pixels and subjected to non-uniform refinement. This yielded consensus map at 2.1 Å resolution (by gold-standard Fourier shell correlation at 0.143 criterion). The resulting map again exhibited reduced resolution at the distal ends of the complex (AccA3 at top and bottom), attributed to conformational flexibility. To further improve the reconstruction, the non-uniform refinement map was used for mask generation. Density subtractions were employed to isolate the AccA3 at top, 4AccD5:2AccD4:2AccE5 at centre, and AccA3 at bottom regions. Focused refinement of the centre region (4AccD5:2AccD4:2AccE5) yielded a locally refined map at 2.1 Å. Given the substantial conformational variability, 3D variability analysis was performed on the 4AccD5:2AccD4:2AccE5 and the resulting principal components were analyzed in ChimeraX v1.9. Based on the principal component analysis, particles were subjected to 3D variability display (cluster mode) job in cryoSPARC v4.5.3 (Structura Biotechnology Inc.), and the best-resolved classes were subjected to non-uniform refinement, result in two distinct complexes 4AccD5:2AccD4:2AccE5 and 6AccD5:2AccE5. The final reconstruction result in resolution 2.1 Å for 4AccD5:2AccD4:2AccE5 and 2.4 Å for 6AccD5:2AccE5, respectively, which were subsequently used for atomic model building. To resolve flexible AccA3, direct local refinement from particles subtraction of the flexible AccA3 at top and bottom regions did not produce substantial improvements. Then subtracted particles corresponding to these regions were subjected to ab initio reconstruction to generate three initial models. These models were used in heterogeneous refinement to classify the particle population. The best-resolved classes were further subjected to non-uniform refinement, resulting in improved density maps for the AccA3 at top and bottom regions. Final resolutions, determined at the gold-standard FSC 0.143 criterion, were 2.92 Å and 3.01 Å for AccA3 at top and AccA3 at bottom, respectively, and were subsequently used to guide atomic model building. Further, to get full map of both the complexes, 3D classification was performed on full set of particles from non-uniform refinement. Two classes with 161,800 and 105,083 particles were selected where the full reconstruction of LCC (4AccD5:2AccD4:8AccA3:2AccE5) and 8AccA3:6AccD5:2AccE5 complex could be resolved. Further particles from both the classes were subjected to non-uniform refinement, and a similar workflow as above was followed to get high resolution of flexible AccA3 at top and bottom for both the classes. The resulting maps from both the classes were combined to make respective composite maps (Supplementary Fig. 8, 17). These two composite maps result in two clearly distinct complexes, where centre CT module consists of 4AccD5:2AccD4:2AccE5 and 6AccD5:2AccE5.

For the LCC complex incubated with propionyl-CoA and palmitoyl-CoA cryo-EM data (Supplementary Fig. 12), a similar workflow is followed. Particles were extracted with a box size of 480 pixels, Fourier cropped to 220 pixels. Multiple rounds of 2D classification were performed to remove junk particles. An initial ab initio reconstruction was generated from selected particles, followed by iterative heterogeneous refinement to resolve compositional heterogeneity. Particles corresponding to the best-resolved classes were re-extracted with a box size of 480 pixels, Fourier cropped to 416 pixels and subjected to non-uniform refinement. This yielded consensus map at 2.1 Å resolution (by gold-standard Fourier shell correlation at 0.143 criterion). Again, the resulting map exhibited reduced resolution at the distal ends of the complex (AccA3 at top and bottom), attributed to conformational flexibility. To further improve the reconstruction, the non-uniform refinement map was used for mask generation. Density subtractions were employed to isolate the AccA3 at top, 4AccD5:2AccD4:2AccE5 at centre, and AccA3 at bottom regions. Focused refinement of the centre region (4AccD5:2Acc D4:2AccE5) yielded a locally refined map at 2.09 Å. Given the substantial conformational variability, 3D variability analysis was performed on the 4AccD5:2AccD4:2AccE5, and the resulting principal component were analyzed in ChimeraX v1.9. Based on the principal component analysis, particles were subjected to 3D variability display (cluster mode) job in cryoSPARC v4.5.3 (Structura Biotechnology Inc.), and the best-resolved classes were subjected to non-uniform refinement, result in again two distinct complexes 4AccD5:2AccD4:2AccE5 and 6AccD5:2AccE5. The final reconstructions result in 2.2 Å resolution for the 4AccD5:2AccD4:2AccE5 and 2.5 Å for the 6AccD5:2AccE5, respectively (Supplementary Fig. 15a,b), that were subsequently used for atomic model building. To resolve flexible AccA3, again direct local refinement from particles subtraction of the flexible AccA3 at top and bottom regions did not produce substantial improvements. Then subtracted particles corresponding to these regions were subjected to ab initio reconstruction to generate three initial models. These models were used in heterogeneous refinement to classify the particle population. The best-resolved classes were further subjected to non-uniform refinement, resulting in improved density maps for the top and bottom regions. Final resolutions, determined at the gold-standard Fourier shell correlation (FSC) 0.143 criterion, were 2.9 Å, and 3.0 Å respectively (Supplementary Fig. 15c,d), and were subsequently used to guide atomic model building. Further, to get full map of both the complexes, 3D classification was performed on full set of particles from non-uniform refinement. Class with 13,970 particles was selected where full LCC complex (4AccD5:2AccD4:8AccA3:2AccE5) could be resolved. These particles subjected to non-uniform refinement and further local refinement to get high resolution of AccA3 at top, 4AccD5:2AccD4:2AccE5 at centre, and AccA3 at bottom. The resulting locally refined maps were combined to generate a composite reconstruction of the full LCC complex (Supplementary Fig. 11). However, again, we could not resolve the full reconstruction of the other complex.

### Model Building

To model the LCC complex (4AccD5:2AccD4:8AccA3:2AccE5), an initial atomic model of the carboxyl transferase module (4AccD5:2AccD4:2AccE5) was generated using ModelAngelo^34^ and rigid-body fitted into the cryo-EM density in ChimeraX^35^, followed by real-space refinement^36^ in Phenix^37^. All side chains were manually inspected and adjusted, and missing loops and termini were built in Coot^38^ wherever well-defined density was present. Residues lacking interpretable side-chain density were modelled using only the underlying secondary-structure geometry in Coot, and residues with no observable density were either assigned zero occupancy or omitted from the final model. For the flexible biotin carboxylase top and bottom module (4AccA3:1AccE5), individual AlphaFold models were used as starting templates, and each polypeptide chain was independently fitted into the corresponding EM density in ChimeraX. Subsequent real-space refinement was performed in Phenix, followed by manual inspection and adjustment in Coot to refine side chains and to build missing loops and termini where the density allowed reliable interpretation. The B domain (residues 139-210) and BCCP domain (residues 530-598) of AccA3 were included in the models only in regions where the EM density was sufficiently well resolved; in the absence of clear density, these segments were not modelled.

To model the (8AccA3:6AccD5:2AccE5) complex, an AlphaFold-predicted structure was used as the starting template for the carboxyl transferase module (6AccD5:2AccE5) and biotin carboxylase module (4AccA3:1AccE5) and fitted into the corresponding EM density in ChimeraX. Subsequent real-space refinement was performed in Phenix, followed by manual inspection and adjustment in Coot. Residues lacking interpretable side-chain density were modelled using only the underlying secondary-structure geometry in Coot, and residues with no observable density were either assigned zero occupancy or omitted from the final model. All ligand molecules were placed into the EM density using Coot with monomer library restraints and refined in Phenix. To accurately describe the covalent linkage between Lys564 of the BCCP domain and biotin, a covalent bond definition was first introduced in ChimeraX, and a custom restraint describing the Lys-biotin covalent geometry was generated with the elbow in Phenix. This restraint was applied during the final rounds of real-space refinement.

### Liquid chromatography/mass spectrometry

Separation of different chain-length acyl-CoAs was achieved by ion pairing-reverse phase high-performance liquid chromatography (IP-RP-HPLC) with an Agilent 1260 Infinity II instrument (Agilent Corporation, Santa Clara, CA, USA) using a NUCLEODUR EC C18 column (dimensions: 50mm x 4.6mm x 3μm). The flow rate was 1ml/min in gradient mode with following elution program: the column was equilibrated with mobile phase A [90% H_2_O and 10% (v/v) 100mM ammonium acetate]. 10 µl of each sample was injected, and same elution condition continued for 1.5 min, followed by 9 min gradient to mobile phase B [90% CH_3_CN and 10% (v/v) 100mM ammonium acetate]. Afterward, the column was washed for 2 min with mobile phase B [90% CH_3_CN and 10% (v/v) 100mM ammonium acetate], followed by 10 sec gradients to reach mobile phase A [90% H_2_O and 10% (v/v) 100mM ammonium acetate] and holding 2 more minutes at this conditions to equilibrate the column before injection of the next sample.

Mass spectrometry was performed using an Agilent 6475 LC/TQ system (Agilent Technologies, Santa Clara, CA, USA) equipped with an AJS electrospray ionization (ESI) source, operated in positive ion mode. ESI parameters were optimized for acyl-CoA’s detection as follows: Nebulizer pressure 20.0 psi, gas flow 10.0 L/min, dry heater 100 °C, capillary voltage 4,000 V. The target mass scan was set from 100 m/z to 1,500 m/z. The fragmentor voltage and collision energy for the tandem mass spectrometry were individually optimized for each acyl-CoA to maximize signal intensity and fragmentation efficiency.

### *In vitro* LCC activity assay

LCC enzymatic activity was quantified by detecting carboxylated acyl-CoA products using ion-pair reverse-phase high-performance liquid chromatography (HPLC) coupled to triple quadrupole mass spectrometry (TQMS/MS). Reactions were carried out in a total volume of 35 μl containing 50 mM HEPES (pH 7.2), 2 mM ATP, 8 mM MgCl₂, 50 mM NaHCO₃, and 50 μM acyl-CoA substrate. The reactions were initiated by the addition of purified LCC complex (final concentration 0.25 μM) and incubated at 30 °C for 4 h. To quench enzymatic activity, reaction mixtures were transferred to -80 °C, followed by denaturation at 70 °C for 10 min. After centrifugation, the resulting supernatant was collected for further analysis. An internal standard was added prior to injection, and 10 μl of the diluted reaction mixture was loaded onto the chromatographic column. Acyl-CoA species were separated by ion-pairing reverse-phase HPLC. Detection was performed on an Agilent 6475 LC/TQ system (Agilent Technologies, Santa Clara, CA, USA) equipped with an AJS electrospray ionization (ESI) source, operated in positive ion mode. For substrate competition assays, equimolar (50 μM) mixtures of acyl-CoA species with varying carbon chain lengths were included, and reactions were conducted similarly as described above except that reactions were quenched after 10 min.

### Data Analysis

Quantitative analyses were performed as follows. Calibration curves were first generated for individual acyl-CoA standards (C3, C8, C10, C12, C14, and C16-CoA) to confirm linearity between chromatographic peak area and analyte concentration. Extracted-ion chromatogram peaks corresponding to carboxylated coenzyme A, uncarboxylated coenzyme A, and the internal standard were integrated using MassHunter Qualitative Analysis software v10.0 (Agilent Technologies, Santa Clara, CA, USA), and the resulting peak areas were exported to spreadsheets and further processed in GraphPad Prism 10 (GraphPad Software, San Diego, CA). All samples were analyzed in triplicate, and data are reported as mean values ± standard deviation.

### ATPase activity assay

The ATP hydrolysis activity of the LCC complex was measured spectrophotometrically^39^ by coupling ADP production to NADH oxidation using pyruvate kinase and lactate dehydrogenase. Oxidation of NADH was monitored at 340 nm in a microplate reader (FLUOstar Omega, BMG LABTECH). Reactions were carried out in a total volume of 80 μl in 384-well non-binding microplates (Greiner). The assay buffer contained 50 mM HEPES (pH 7.6), 50 mM NaHCO₃, 5 mM MgCl₂, 3 mM ATP, 0.5 mM phosphoenolpyruvate, 0.2 mM NADH, 0.3 mg/ml BSA, 4.8-8 units of pyruvate kinase, 7.2-11.2 units of lactate dehydrogenase, and the desired concentration of acyl-CoA. Reactions were initiated by addition of purified LCC complex to a final concentration of 0.125 μM. Assays were performed at 30 °C using preincubated plates.

### Structure predictions

The structures of the 4AccD5:2AccD4:2AccE5 and 4AccA3:1AccE5 complexes from different *Mycobacteria* and *Corynebacteria* species were predicted using AlphaFold3^40^ (DeepMind, 2025 release). Protein sequences were retrieved from UniProt and submitted to AlphaFold3 with the appropriate stoichiometry and default settings. For each complex, among the five structural models generated, the model with the highest confidence was selected. The structures were visualized with UCSF ChimeraX 1.9^35^. Subunits were coloured according to the proposed scheme, and for the AccD4/AccD5/AccE5 complex the unstructured N-terminal region of AccE5 was removed for clarity.

## Data and Material Availability

Cryo-EM maps and associated atomic models reported in this study have been deposited in the Electron Microscopy Data Bank under accession codes EMD-56274 (8AccA3:6AccD5:2AccE5 complex bound to ADP, and Mg_2+_, non-uniform refinement (consensus map)), EMD-56275 (8AccA3:6AccD5:2AccE5 complex bound to ADP, and Mg_2+_, non-uniform refinement (focused map of bottom BC module and centre CT module)), EMD-56276 (8AccA3:6AccD5:2AccE5 complex bound to ADP, and Mg_2+_, non-uniform refinement (focused map of top BC module and centre CT module)), EMD-56277 (8AccA3:6AccD5:2AccE5 complex bound to ADP, and Mg_2+_, local refinement (focused map of centre CT module), EMD-56278 (8AccA3:6AccD5:2AccE5 complex bound to ADP, and Mg_2+_, non-uniform refinement (focused map of bottom BC module)), EMD-56279 (LCC complex bound to ADP, and Mg_2+_, non-uniform refinement (focused map of top BC module)), **EMD-56280** (8AccA3:6AccD5:2AccE5 complex bound to ADP, and Mg_2+_, composite map), EMD-56287 (LCC complex bound to palmitoyl-CoA, ADP, and Mg_2+_, non-uniform refinement (consensus map)), EMD-56288 (LCC complex bound to palmitoyl-CoA, ADP, and Mg_2+_, non-uniform refinement (focused map of top BC module and centre CT module)), EMD-56289 (LCC complex bound to palmitoyl-CoA, ADP, and Mg_2+_, non-uniform refinement (focused map of bottom BC module and centre CT module)), EMD-56290 (LCC complex bound to palmitoyl-CoA, ADP, and Mg_2+_, non-uniform refinement (focused map of top BC module)), EMD-56291 (LCC complex bound to palmitoyl-CoA, ADP, and Mg_2+_, non-uniform refinement (focused map of bottom BC module)), EMD-56292 (LCC complex bound to palmitoyl-CoA, ADP, and Mg_2+_, local refinement (focused map of centre CT module)), **EMD-56293** (LCC complex bound to palmitoyl-CoA, ADP, and Mg_2+_, composite map), **EMD-56369** (BC top module (4AccA3:1AccE5) bound to ADP, and Mg_2+_, non-uniform refinement of full data set), **EMD-56370** (BC bottom module (4AccA3:1AccE5) bound to ADP, and Mg_2+_, local refinement of full data set), EMD-56301 (LCC complex bound to palmitoyl-CoA, non-uniform refinement (consensus map)), EMD-56302 (LCC complex bound to palmitoyl-CoA, non-uniform refinement (focused map of top BC module and centre CT module)), EMD-56303 (LCC complex bound to palmitoyl-CoA, non-uniform refinement (focused map of bottom BC module and centre CT module)), EMD-56304 (LCC complex bound to palmitoyl-CoA, local refinement (focused map of centre CT module)), EMD-56305 (LCC complex bound to palmitoyl-CoA, local refinement (focused map of top BC module sub volume 1)), EMD-56306 (LCC complex bound to palmitoyl-CoA, local refinement (focused map of top BC module sub volume 2)), EMD-56307 (LCC complex bound to palmitoyl-CoA, local refinement (focused map of bottom BC module sub volume 1)), EMD-56308 (LCC complex bound to palmitoyl-CoA, local refinement (focused map of bottom BC module sub volume 2)), **EMD-56309** (LCC complex bound to palmitoyl-CoA, non-uniform refinement (composite map)), **EMD-56336** (Centre CT module (4AccD5:2AccD4:2AccE5) bound to palmitoyl-CoA, non-uniform refinement of cluster 1 from 3D variability analysis), **EMD-56359** (Centre CT module (6AccD5), non-uniform refinement of cluster 2 from 3D variability analysis), **EMD-56360** (BC top module (4AccA3:1AccE5), non-uniform refinement of full data set), **EMD-56362** (BC bottom module (4AccA3:1AccE5), non-uniform refinement of full data set), EMD-56314 (LCC complex bound to palmitoyl-CoA and propionyl-CoA, non-uniform refinement (consensus map)), EMD-56315 (LCC complex bound to palmitoyl-CoA and propionyl-CoA, non-uniform refinement (focused map of top BC module and centre CT module)), EMD-56317 (LCC complex bound to palmitoyl-CoA and propionyl-CoA, non-uniform refinement (focused map of bottom BC module and centre CT module)), EMD-56318 (LCC complex bound to palmitoyl-CoA and propionyl-CoA, local refinement (focused map of centre CT module)), EMD-56320 (LCC complex bound to palmitoyl-CoA and propionyl-CoA, local refinement (focused map of top BC module)), EMD-56321 (LCC complex bound to palmitoyl-CoA and propionyl-CoA, local refinement (focused map of bottom BC module)), **EMD-56323** (LCC complex bound to palmitoyl-CoA and propionyl-CoA, composite map), **EMD-56332** (Centre CT module (4AccD5:2AccD4:2AccE5) bound to palmitoyl-CoA and propionyl-CoA, non-uniform refinement of cluster 1 from 3D variability analysis), **EMD-56333** (Centre CT module (6AccD5:2AccE5) bound to propionyl-CoA, non-uniform refinement of cluster 2 from 3D variability analysis), **EMD-56334** (BC top module (4AccA3:1AccE5), non-uniform refinement of full data set), **EMD-56335** (BC bottom module (4AccA3:1AccE5), non-uniform refinement of full data set); and the Protein Data Bank with accession codes **PBD_00009TUV** (8AccA3:6AccD5:2AccE5 complex bound to ADP, and Mg_2+_), **PDB_00009TV3** (LCC complex bound to palmitoyl-CoA, ADP, and Mg_2+_), **PBD_00009TWN** (BC top module (4AccA3:1AccE5) bound to ADP, and Mg_2+_), **PBD_00009TWO** (BC bottom module(4AccA3:1AccE5) bound to ADP, and Mg_2+_), **PBD_00009TV6** (LCC complex bound to palmitoyl-CoA), **PBD_00009TVS** (Centre CT module (4AccD5:2AccD4:2AccE5) bound to palmitoyl-CoA), **PBD_00009TW5** (Ce ntre CT module (6AccD5)), **PBD_00009TW6** (BC top module (4AccA3:1AccE5) from EM data of LCC complex incubated with palmitoyl-CoA)), **PBD_00009TWI** (BC bottom module (4AccA3:1AccE5) from EM data of LCC complex incubated with palmitoyl-CoA EM data), **PBD_00009TVE** (LCC complex bound to palmitoyl-CoA and propionyl-CoA), **PBD_00009TVM** (Centre CT module (4AccD5:2AccD4:2AccE5) bound to palmitoyl CoA and propionylCoA), **PBD_00009TVO** (Centre CT module (6AccD5:2AccE5) bound to propionyl CoA), **PBD_00009T VP** (BC top module (4AccA3:1AccE5) from EM data of LCC complex incubated with palmitoyl-CoA and propionyl-CoA), **PBD_00009TVQ** (BC bottom module (4AccA3:1AccE5) from EM data of LCC complex incubated with palmitoyl-CoA and propionyl-CoA).

## Acknowledgments

We thank Hans van den Elst for assistance with mass spectrometry, Patrick Voskamp for laboratory support, Kanwal Kayastha for advice on data processing at Leiden Institute of Chemistry. We also thank Willem Noteborn and Ludovic Renault (Netherlands Centre for Electron Nanoscopy) for support with cryo-EM data collection.

This work was supported by the Leiden Institute of Chemistry and by Oncode Accelerator, a Dutch National Growth Fund project under grant number NGFPO2201. It benefited from access to the Netherlands Centre for Electron Nanoscopy (NeCEN) as part of the Netherlands Electron Microscopy Infrastructure (project no. 184.034.014) of the National Roadmap for Large-Scale Research Infrastructure of the Dutch Research Council.

## Author contributions

A.Y. and S.G. designed research; A.Y. and B.F. performed research; A.Y. performed the biochemistry, sample preparation, grid screening, image processing and reconstructions and build the model; B.F. performed LC MS, A.Y., N.R., B.F., S.G. analyzed data; A.Y., N.R.,B.F. and S.G. wrote the paper. The authors declare no competing interest.

## Notes

### Competing Interest Statement

The authors have declared no competing interest.

### Summary of Updates

Version update statement In the course of preparing cryo-EM validation reports for journal submission, we re-evaluated the corresponding reconstructions and map assignments. This analysis revealed that a conformational state previously interpreted as an open form of LCC instead corresponds to an AccD5-centered assembly (ACCase 5). The manuscript has been revised accordingly to correct the structural assignment and refine the mechanistic interpretation. The updated version clarifies the distinction between the AccD4/AccD5 heterohexamer (LCC) and the AccD5 homohexamer (ACCase 5).

