## Supplementary material for "Structural Basis of Substrate Selectivity and Catalysis in the Mycobacterial Long-Chain Acyl-CoA Carboxylase": SUPPLEMENTARY_INFORMATION_submission.pdf

**Supplementary Table 1.** Data collection and reconstruction statistics for the palmitoyl-CoA–bound LCC complex.

| **Data collection** | | | | | | | | |
|---|---|---|---|---|---|---|---|---|
| Magnification | 105000 | | | | | | | |
| Voltage (kV) | 300 | | | | | | | |
| Electron exposure (e-/ Å$^2$) | 60 | | | | | | | |
| Defocus range (μm) | 0.8-2.0 | | | | | | | |
| Raw pixel size (Å) | 0.84 | | | | | | | |
| Micrograph collected | 14,716 | | | | | | | |
| **Reconstruction** | | | | | | | | |
| Symmetry | C1 | | | | | | | |
| Initial particle images extracted (no.) | 1,900,567 | | | | | | | |
| Initial particle images for 3D processing (no.) | 1,252,663 | | | | | | | |
| | Consensus map (EMD-56301) | Focused map of top BC and central CT module (EMD-56302) | Focused map of bottom BC and central CT module (EMD-56303) | Focused map of central CT module (EMD-56304) | Focused map of top BC module sub - volume1 (EMD-56305) | Focused map of top BC sub-volume 2 (EMD-56306) | Focused map of bottom BC module sub - volume1 (EMD-56307) | Focused map of bottom BC module sub - volume2 (EMD-56308) |
| Final Particle images(no.) | 65445 | | | | | | | |
| Map resolution (Å), FSC threshold (0.143) | 2.51 | 2.45 | 2.51 | 2.45 | 3.21 | 3.54 | 3.11 | 3.24 |
| FSC mask map resolution range (Å), FSC threshold(0.143) | 2.5-3.6 | 2.5-3.6 | 2.5-3.6 | 2.5-3.9 | 3.2-6.4 | 3.5-6.8 | 3.1-6.4 | 3.2-6.3 |
| Total map resolution range (Å) | 2.5-40.8 | 2.5-41.1 | 2.5-41.0 | 2.5-41.1 | 3.2-46.2 | 3.5-47.5 | 3.1-46.1 | 3.2-31.0 |
| Map sharpening B- factor | 46.2 | 40.7 | 42.4 | 43.3 | 68.8 | 71.5 | 70.5 | 74.2 |

**Supplementary Table 2.** Data collection and reconstruction statistics for the CT and BC modules from the cryo-EM dataset of the palmitoyl-CoA–bound LCC complex.

| **Data collection** | | | | |
|---|---|---|---|---|
| Magnification | 105000 | | | |
| Voltage (kV) | 300 | | | |
| Electron exposure (e-/ Å$^2$) | 60 | | | |
| Defocus range (μm) | 0.8-2.0 | | | |
| Raw pixel size (Å) | 0.84 | | | |
| Micrograph collected | 14,716 | | | |
| **Reconstruction** | | | | |
| Symmetry | C1 | | | |
| Initial particle images extracted (no.) | 1,900,567 | | | |
| Initial particle images for 3D processing (no.) | 1,252,663 | | | |
| | Central CT module (4AccD5:2AccD4:2AccE5), non-uniform refinement of cluster 1 from 3D variability analysis (EMD-56336) | Central CT module (6AccD5), non-uniform refinement of cluster 2 from 3D variability analysis (EMD-56359) | Top BC module (4AccA3:1AccE5), non-uniform refinement of full data set (EMD-56360) | Bottom BC module (4AccA3:1AccE5), non-uniform refinement of full data set (EMD-56362) |
| Final Particle images(no.) | 156,626 | 19,395 | 324,580 | 326,828 |
| Map resolution (Å), FSC threshold (0.143) | 2.3 | 2.9 | 3.0 | 3.0 |
| FSC mask map resolution range (Å), FSC threshold(0.143) | 2.3-3.2 | 2.9-6.1 | 3.0-3.7 | 3.0-3.7 |
| Total map resolution range (Å) | 2.3-35.3 | 2.9-46.0 | 3.0-51.0 | 3.0-45.5 |
| Map sharpening B- factor | 48.4 | 39.8 | 84.5 | 86.4 |

**Supplementary Table 3.** Atomic model refinement and validation statistics for the palmitoyl-CoA–bound LCC complex

| Model Building and Refinement | | | | | |
|---|---|---|---|---|---|
| | LCC complex bound to palmitoyl-CoA (9TV6, EMD-56293) | Central CT module (4AccD5:2AccD4: 2AccE5) bound to palmitoyl-CoA (9TVS, EMD-56336) | Central CT module (6AccD5) (9TW5, EMD-56359) | Top BC module (4AccA3:1AccE5) (9TW6, EMD-56360) | Bottom BC module (4AccA3: 1AccE5) (9TWI, EMD-56362) |
| Initial model used | Model Angelo, AlphaFold | Model Angelo | AlphaFold | AlphaFold | AlphaFold |
| Model resolution (Å) | 3.1 | 2.6 | 3.1 | 3.4 | 3.3 |
| FSC threshold | 0.5 | 0.5 | 0.5 | 0.5 | 0.5 |
| CC (volume/mask) | 0.82/0.81 | 0.87/0.87 | 0.81/0.83 | 0.83/0.82 | 0.86/0.86 |
| Protein B factor (Å$^2$) | 62.82 | 80.55 | 67.19 | 145.91 | 114.17 |
| Ligand B factor (Å$^2$) | 50.27 | 121.64 | | 137.97 | 20.00 |
| | **Model composition** | | | | |
| Nonhydrogen atom | 52456 | 26471 | 23577 | 14737 | 13858 |
| Protein residue | 6889 | 3459 | 3106 | 1954 | 1825 |
| Ligands | PKZ: 2, BTN: 6 | PKZ: 2, BTN: 4 | | BTN: 4 | BTN: 4 |
| | **RMS deviations** | | | | |
| Bond length (Å) | 0.003 | 0.003 | 0.003 | 0.003 | 0.003 |
| Bond angle (°) | 0.469 | 0.534 | 0.557 | 0.535 | 0.520 |
| | **Validation** | | | | |
| MolProbity score | 1.64 | 1.64 | 1.95 | 1.58 | 2.02 |
| Clashscore | 4.64 | 5.91 | 6.56 | 4.53 | 4.60 |
| Poor rotamers (%) | 2.33 | 1.34 | 2.65 | 2.23 | 4.18 |
| CaBLAM outlier(%) | 1.57 | 1.71 | 1.90 | 1.37 | 2.23 |
| | **Ramachandran plot** | | | | |
| Favored (%) | 97.37 | 96.56 | 96.07 | 97.56 | 95.52 |
| Allowed (%) | 2.62 | 3.30 | 3.60 | 2.44 | 4.36 |
| Disallowed (%) | 0.01 | 0.15 | 0.32 | 0.00 | 0.11 |

**Supplementary Table 4.** Data collection and reconstruction statistics for the palmitoyl-CoA–, $Mg^{2+}$-, and ADP-bound LCC complex and BC modules.

| **Data collection** | | | | | | | | |
|---|---|---|---|---|---|---|---|---|
| Magnification | 105000 | | | | | | | |
| Voltage (kV) | 300 | | | | | | | |
| Electron exposure (e-/ Å$^2$) | 60 | | | | | | | |
| Defocus range (μm) | 0.8-2.0 | | | | | | | |
| Raw pixel size (Å) | 0.84 | | | | | | | |
| Micrograph collected | 15130 | | | | | | | |
| **Reconstruction** | | | | | | | | |
| Symmetry | C1 | | | | | | | |
| Initial particle images extracted (no.) | 3,193,269 | | | | | | | |
| Initial particle images for 3D processing (no.) | 1,974,128 | | | | | | | |
| | Consensus map (EMD-56287) | Focused map of top BC and central CT module (EMD-56288) | Focused map of bottom BC and central CT module (EMD-56289) | Focused map of central CT module (EMD-56292) | Focused map of top BC module (EMD-56290) | Focused map of bottom BC (EMD-56291) | Top BC module (full dataset) (EMD-56369) | Bottom BC module (full dataset) (EMD-56370) |
| Final Particle images(no.) | 161800 | | | | 93766 | 95749 | 210,927 | 194,783 |
| Map resolution (Å), FSC threshold (0.143) | 2.25 | 2.24 | 2.26 | 2.2 | 3.22 | 3.21 | 2.97 | 2.99 |
| FSC mask map resolution range (Å), FSC threshold(0.143) | 2.3-3.1 | 2.2-3.1 | 2.3-3.1 | 2.2-3.4 | 3.2-6.2 | 3.2-6.3 | 3.0-3.8 | 3.0-4.2 |
| Total map resolution range (Å) | 2.3-35.9 | 2.2-35.9 | 2.3-35.8 | 2.2-35.9 | 3.2-50.4 | 3.2-50.5 | 3.0-45.9 | 3.0-47.6 |
| Map sharpening B-factor | 46.7 | 39.0 | 40.8 | 40.4 | 66.3 | 67.2 | 79.3 | 74.4 |

**Supplementary Table 5.** Data collection and reconstruction statistics for the $Mg^{2+}$- and ADP-bound 8AccA3:6AccD5:2AccE5 complex.

| | | | | | | |
|---|---|---|---|---|---|---|
| **Data collection** | | | | | | |
| Magnification | 105000 | | | | | |
| Voltage (kV) | 300 | | | | | |
| Electron exposure (e-/ Å$^2$) | 60 | | | | | |
| Defocus range (µm) | 0.8-2.0 | | | | | |
| Raw pixel size (Å) | 0.84 | | | | | |
| Micrograph collected | 15130 | | | | | |
| **Reconstruction** | | | | | | |
| Symmetry | C1 | | | | | |
| Initial particle images extracted (no.) | 3,193,269 | | | | | |
| Initial particle images for 3D processing (no.) | 1,974,128 | | | | | |
| | Consensus map (EMD-56274) | Focused map of top BC and central CT module (EMD-56275) | Focused map of bottom BC and central CT module (EMD-56276) | Focused map of central CT module (EMD-56277) | Focused map of top BC module (EMD-56279) | Focused map of bottom BC (EMD-56278) |
| Final Particle images(no.) | 105,083 | | | | 57,102 | 55,619 |
| Map resolution (Å), FSC threshold(0.143) | 2.66 | 2.63 | 2.65 | 2.62 | 3.45 | 3.51 |
| FSC mask map resolution range (Å), FSC threshold(0.143) | 2.6-3.5 | 2.6-3.5 | 2.6-3.5 | 2.6-3.9 | 3.4-7.0 | 3.5-7.0 |
| Total map resolution range (Å) | 2.6-41.0 | 2.6-41.0 | 2.6-41.1 | 2.6-41.1 | 3.4-54.3 | 3.5-55 |
| Map sharpening B-factor | 55.6 | 49.5 | 52.1 | 50.0 | 68.3 | 68.2 |

**Supplementary Table 6.** Atomic model refinement and validation statistics for the palmitoyl-CoA-, ATP-, $Mg^{2+}$-, and $NaHCO_3$–bound LCC and 8AccA3:6AccD5:2AccE5 complex.

| Model Building and Refinement | | | | |
|---|---|---|---|---|
| | LCC complex bound to palmitoyl-CoA, $Mg^{2+}$, and ADP (9TV3, EMD-56293) | 8AccA3:6AccD5:2AccE5 complex bound to $Mg^{2+}$, and ADP (9TUV, EMD-56280) | Top BC module (4AccA3:1AccE5) (9TWN, EMD-56369) | Bottom BC module (4AccA3:1AccE5) (9TWO, EMD-56370) |
| Initial model used | Model Angelo, AlphaFold | Model Angelo, AlphaFold | AlphaFold | AlphaFold |
| Model resolution (Å) | 2.3 | 2.8 | 3.5 | 3.4 |
| FSC threshold | 0.5 | 0.5 | 0.5 | 0.5 |
| CC (volume/mask) | 0.87/0.86 | 0.79/0.79 | 0.83/0.83 | 0.82/0.82 |
| Protein B factor (Å$^2$) | 63.89 | 80.24 | 124.86 | 133.91 |
| Ligand B factor (Å$^2$) | 49.16 | 129.19 | 20.0 | 20.0 |
| **Model composition** | | | | |
| Nonhydrogen atom | 51657 | 51406 | 13998 | 13952 |
| Protein residue | 6742 | 6733 | 1830 | 1823 |
| Ligands | PKZ:2, MG:7, ADP:8 | MG:4, ADP:8 | MG:4, ADP:4 | MG:4, ADP:4 |
| **RMS deviations** | | | | |
| Bond length (Å) | 0.003 | 0.004 | 0.004 | 0.004 |
| Bond angle (°) | 0.619 | 0.655 | 0.671 | 0.741 |
| **Validation** | | | | |
| MolProbity score | 1.84 | 2.06 | 1.96 | 2.14 |
| Clashscore | 5.61 | 7.19 | 5.98 | 7.15 |
| Poor rotamers (%) | 2.52 | 3.51 | 3.34 | 4.62 |
| CaBLAM outlier(%) | 2.08 | 2.24 | 2.40 | 2.24 |
| **Ramachandran plot** | | | | |
| Favored (%) | 96.49 | 96.31 | 96.47 | 96.34 |
| Allowed (%) | 3.35 | 3.49 | 3.15 | 3.49 |
| Disallowed (%) | 0.17 | 0.19 | 0.39 | 0.17 |

**Supplementary Table 7.** Data collection and reconstruction statistics for the palmitoyl-CoA– and propionyl-CoA–bound LCC complex.

| **Data collection** | | | | | | |
|---|---|---|---|---|---|---|
| Magnification | 105000 | | | | | |
| Voltage (kV) | 300 | | | | | |
| Electron exposure (e-/ Å$^2$) | 60 | | | | | |
| Defocus range (μm) | 0.8-2.0 | | | | | |
| Raw pixel size (Å) | 0.84 | | | | | |
| Micrograph collected | 14,974 | | | | | |
| **Reconstruction** | | | | | | |
| Symmetry | C1 | | | | | |
| Initial particle images extracted (no.) | 3,308,143 | | | | | |
| Initial particle images for 3D processing (no.) | 1,707,887 | | | | | |
| | Consensus map (EMD-56314) | Focused map of top BC and central CT module (EMD-56315) | Focused map of bottom BC and central CT module (EMD-56317) | Focused map of central CT module (EMD-56318) | Focused map of top BC module (EMD-56320) | Focused map of bottom BC module (EMD-56321) |
| Final Particle images(no.) | 13970 | | | | | |
| Map resolution (Å), FSC threshold (0.143) | 2.9 | 2.8 | 2.8 | 2.8 | 4.1 | 4.4 |
| FSC mask map resolution range (Å), FSC threshold(0.143) | 2.9-6.9 | 2.8-6.9 | 2.8-6.9 | 2.8-7.1 | 4.1-8.5 | 4.4-8.5 |
| Total map resolution range (Å) | 2.9-51.1 | 2.8-46.3 | 2.8-46.2 | 2.8-46.2 | 4.1-62.2 | 4.4-60.7 |
| Map sharpening B-factor | 24.7 | 24.5 | 25.4 | 28.3 | 61.3 | 68.2 |

**Supplementary Table 8.** Data collection and reconstruction statistics for the CT and BC modules from the cryo-EM dataset of the palmitoyl-CoA– and propionyl-CoA–bound LCC complex.

| **Data collection** | | | | |
|---|---|---|---|---|
| Magnification | 105000 | | | |
| Voltage (kV) | 300 | | | |
| Electron exposure (e-/ Å$^2$) | 60 | | | |
| Defocus range (μm) | 0.8-2.0 | | | |
| Raw pixel size (Å) | 0.84 | | | |
| Micrograph collected | 14,974 | | | |
| **Reconstruction** | | | | |
| Symmetry | C1 | | | |
| Initial particle images extracted (no.) | 3,308,143 | | | |
| Initial particle images for 3D processing (no.) | 1,707,887 | | | |
| | Central CT module (4AccD5:2AccD4:2AccE5), non-uniform refinement of cluster 1 from 3D variability analysis (EMD-56332) | Central CT module (6AccD5:2AccE5), non-uniform refinement of cluster 2 from 3D variability analysis (EMD-56333) | Top BC module (4AccA3:1AccE5), non-uniform refinement of full data set (EMD-56334) | Bottom BC module (4AccA3:1AccE5), non-uniform refinement of full data set (EMD-56335) |
| Final Particle images(no.) | 244,412 | 195,939 | 436,553 | 425,573 |
| Map resolution (Å), FSC threshold (0.143) | 2.2 | 2.5 | 2.9 | 2.9 |
| FSC mask map resolution range (Å), FSC threshold(0.143) | 2.2-3.0 | 2.5-3.3 | 2.9-3.7 | 2.9-3.7 |
| Total map resolution range (Å) | 2.2-35.5 | 2.5-39.9 | 2.9-28.1 | 2.9-33.4 |
| Map sharpening B- factor | 47.3 | 54.9 | 97.2 | 92.4 |

**Supplementary Table 9.** Atomic model refinement and validation statistics for the palmitoyl-CoA– and propionyl-CoA–bound LCC complex.

| Model Building and Refinement | | | | | |
|---|---|---|---|---|---|
| | LCC complex bound to palmitoyl-CoA and propionyl -CoA (9TVE, EMD-56323) | Central CT module (4AccD5:2AccD4:2 AccE5) bound to palmitoyl-CoA and propionyl-CoA (9TVM, EMD-56332) | Central CT module (6AccD5: 2AccE5) bound to palmit oyl-CoA (9TVO, EMD-56333) | Top BC module (4AccA3:1Acc E5) (9TVP, EMD-56334) | Bottom BC module (4Acc A3:1AccE5) (9TVQ, EMD-56335) |
| Initial model used | Model Angelo, AlphaFold | Model Angelo | AlphaFold | AlphaFold | AlphaFold |
| Model resolution (Å) | 3.6 | 2.5 | 2.8 | 3.3 | 3.3 |
| FSC threshold | 0.5 | 0.5 | 0.5 | 0.5 | 0.5 |
| CC (volume/mask) | 0.80/0.78 | 0.89/0.89 | 0.81/0.81 | 0.84/0.84 | 0.86/0.86 |
| Protein B factor ($Å^2$) | 119.91 | 71.37 | 104.41 | 145.28 | 134.14 |
| Ligand B factor ($Å^2$) | 45.49 | 47.79 | 153.83 | 133.17 | 121.89 |
| | **Model composition** | | | | |
| Nonhydrogen atom | 53105 | 26707 | 24884 | 14774 | 14705 |
| Protein residue | 6952 | 3463 | 3231 | 1956 | 1948 |
| Ligands | PKZ: 2, 1VU: 4 BTN: 6 | PKZ: 2, 1VU: 4 BTN: 4 | 1VU: 6 | BTN: 4 | BTN: 4 |
| | **RMS deviations** | | | | |
| Bond length (Å) | 0.003 | 0.002 | 0.004 | 0.006 | 0.003 |
| Bond angle (°) | 0.520 | 0.471 | 0.526 | 0.546 | 0.492 |
| | **Validation** | | | | |
| MolProbity score | 1.76 | 1.57 | 1.74 | 1.53 | 1.61 |
| Clashscore | 6.95 | 3.77 | 5.44 | 4.52 | 4.74 |
| Poor rotamers (%) | 1.38 | 1.48 | 2.19 | 1.83 | 3.02 |
| CaBLAM outlier(%) | 2.75 | 1.94 | 2.03 | 1.74 | 1.32 |
| | **Ramachandran plot** | | | | |
| Favored (%) | 96.00 | 96.04 | 96.92 | 97.51 | 98.33 |
| Allowed (%) | 3.78 | 3.76 | 2.83 | 2.44 | 1.62 |
| Disallowed (%) | 0.22 | 0.20 | 0.25 | 0.05 | 0.05 |

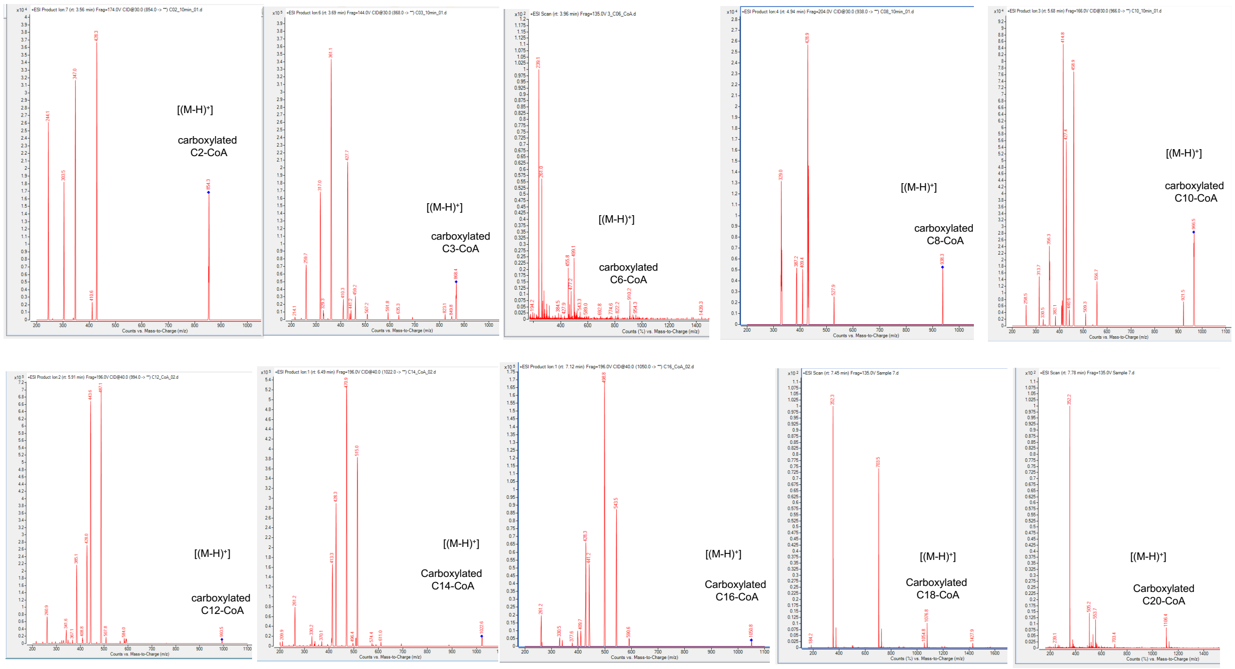


**Supplementary Fig. 1.** Fragmentation patterns of carboxylated acyl-CoA species analyzed by IP-RP-HPLC–ESI–TQMS. Product ion (MS/MS) scans were used for quantification of C2-, C3-, C8-, C10-, C12-, C14-, and C16-CoA, whereas C6-, C18-, and C20-CoA were detected using single-stage mass spectrometry (MS).

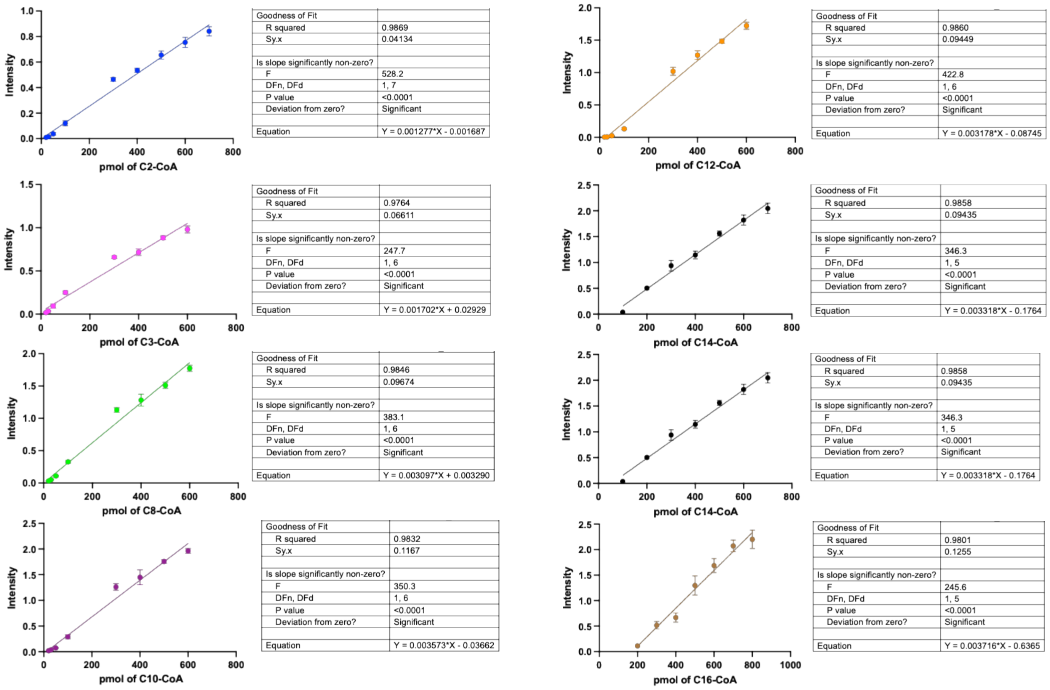


**Supplementary Fig. 2.** Calibration curves for acyl-CoA signal intensities measured by IP-RP-HPLC–ESI–TQMS.

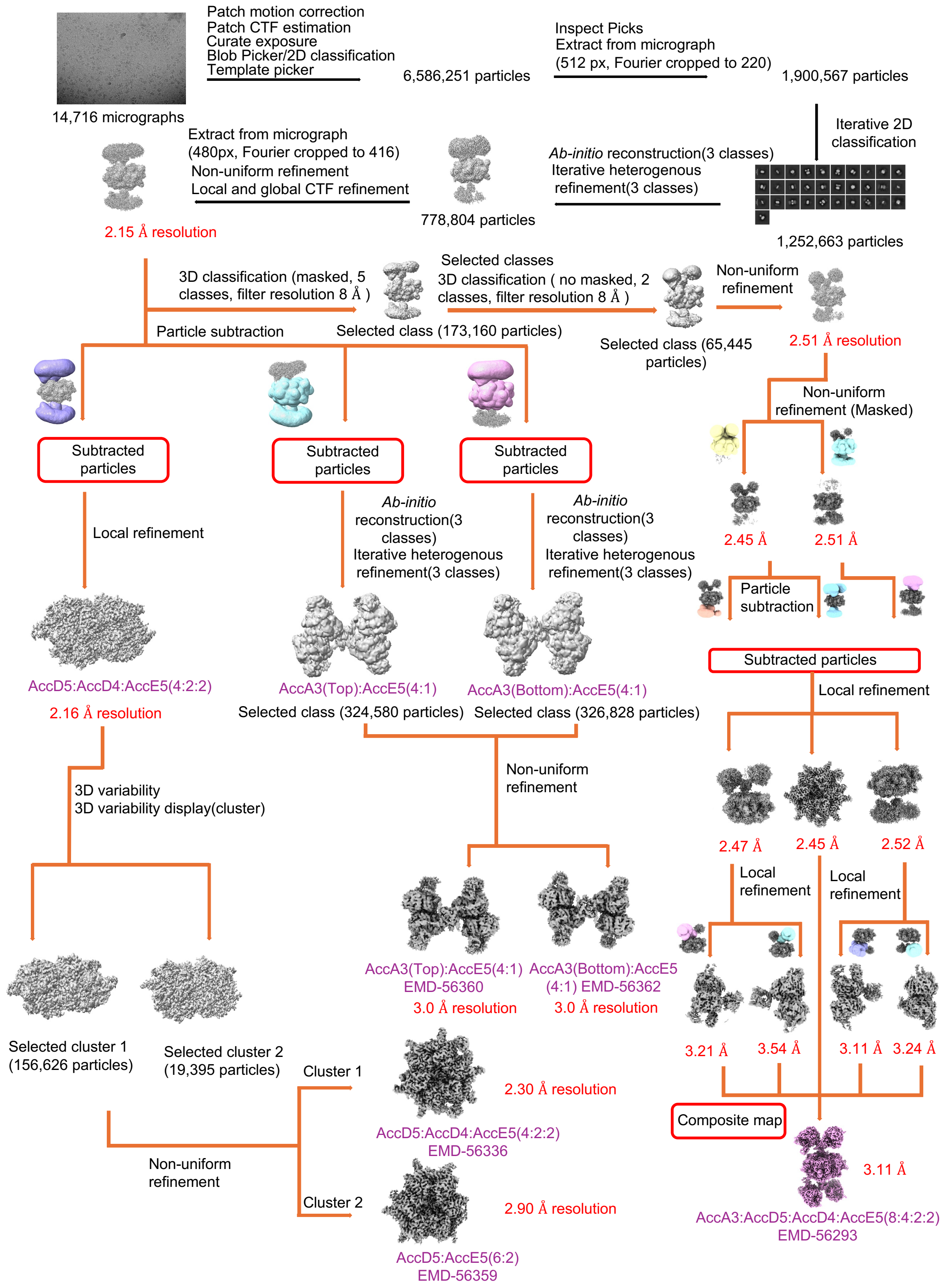


**Supplementary Fig. 3.** Image processing and classification workflow for the cryo-EM dataset of the palmitoyl-CoA–bound LCC complex.

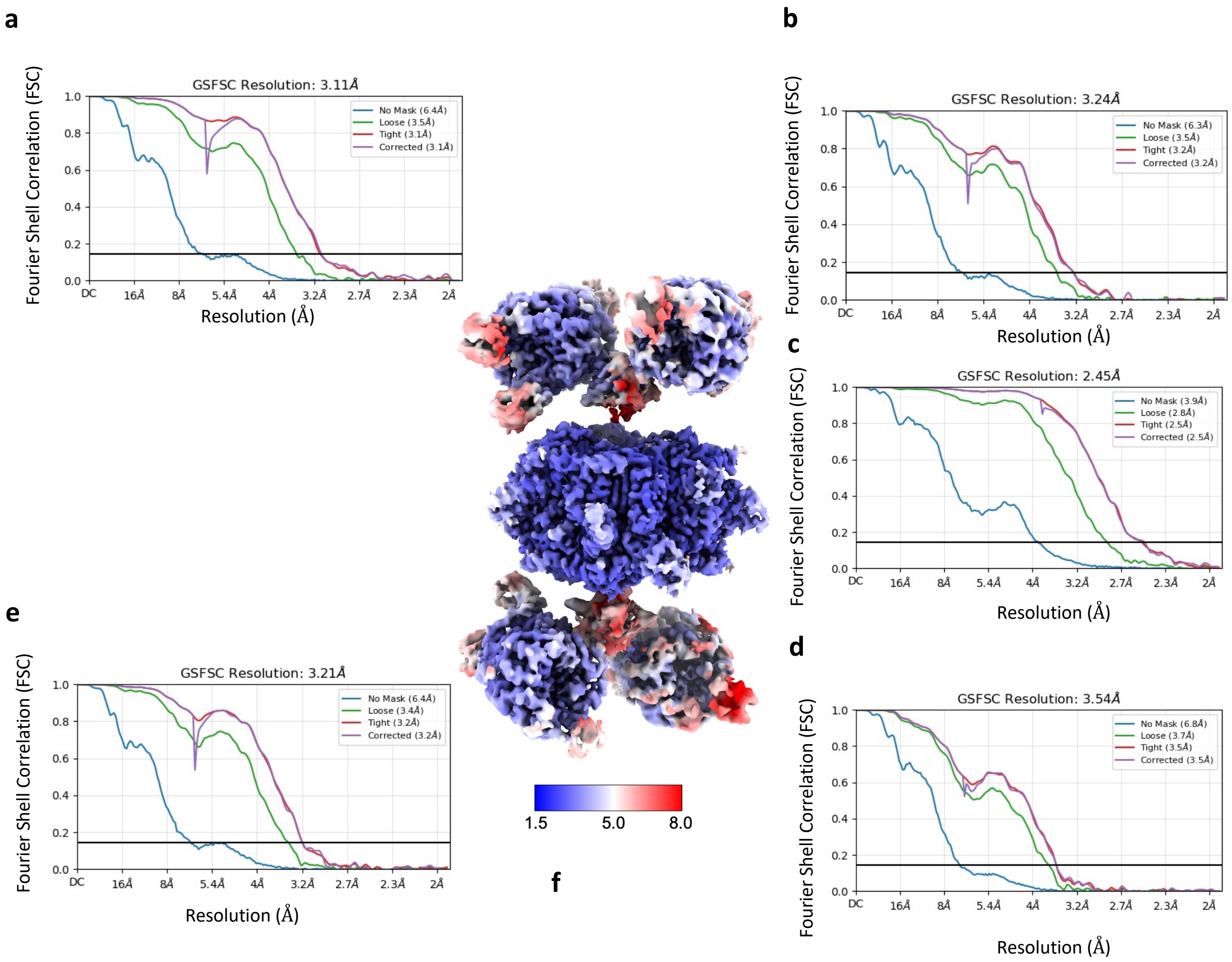


**Supplementary Fig. 4**. Fourier shell correlation (FSC) curves for focused reconstructions of the palmitoyl-CoA–bound LCC complex: **a,** top BC module, subvolume 1; **b,** top BC module, subvolume 2; **c,** central CT module; **d,** bottom BC module, subvolume 2; and **e,** bottom BC module, subvolume 1. **f,** Estimated average resolution of the composite cryo-EM map.

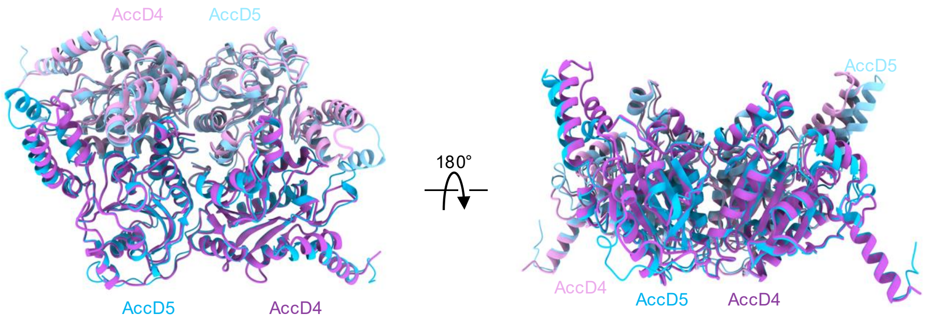


**Supplementary Fig. 5.** Structural superposition of the AccD4 dimer (purple) and the AccD5 dimer (cornflower blue), highlighting their high degree of structural similarity.

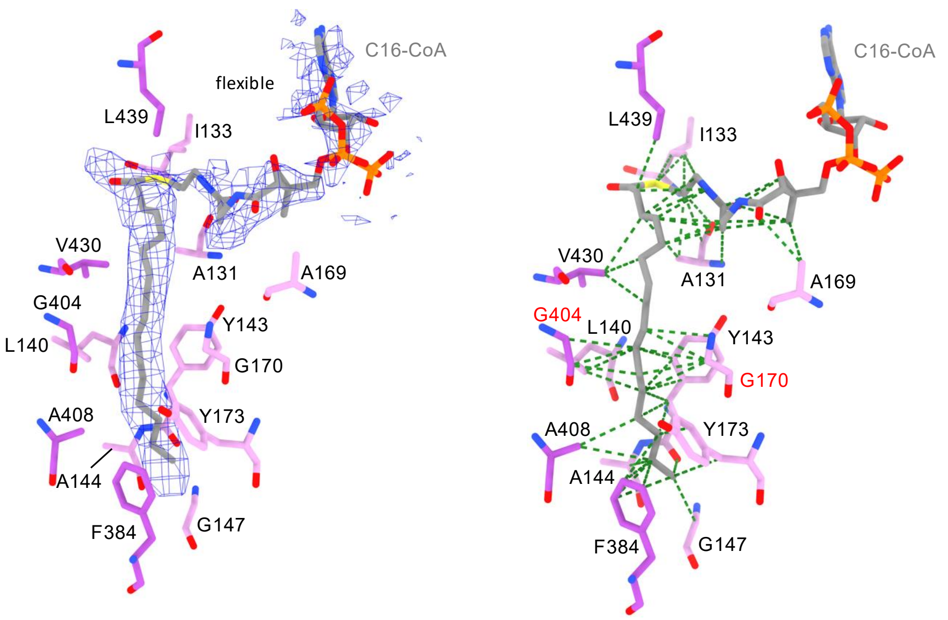


**Supplementary Fig. 6.** Representative density for C16-CoA (blue mesh) bound in the second active site of AccD4 (left). The acyl chain is well resolved and primarily stabilized by van der Waals interactions (right, green dashed lines), whereas the density for the CoA moiety is fragmented indicating flexibility.

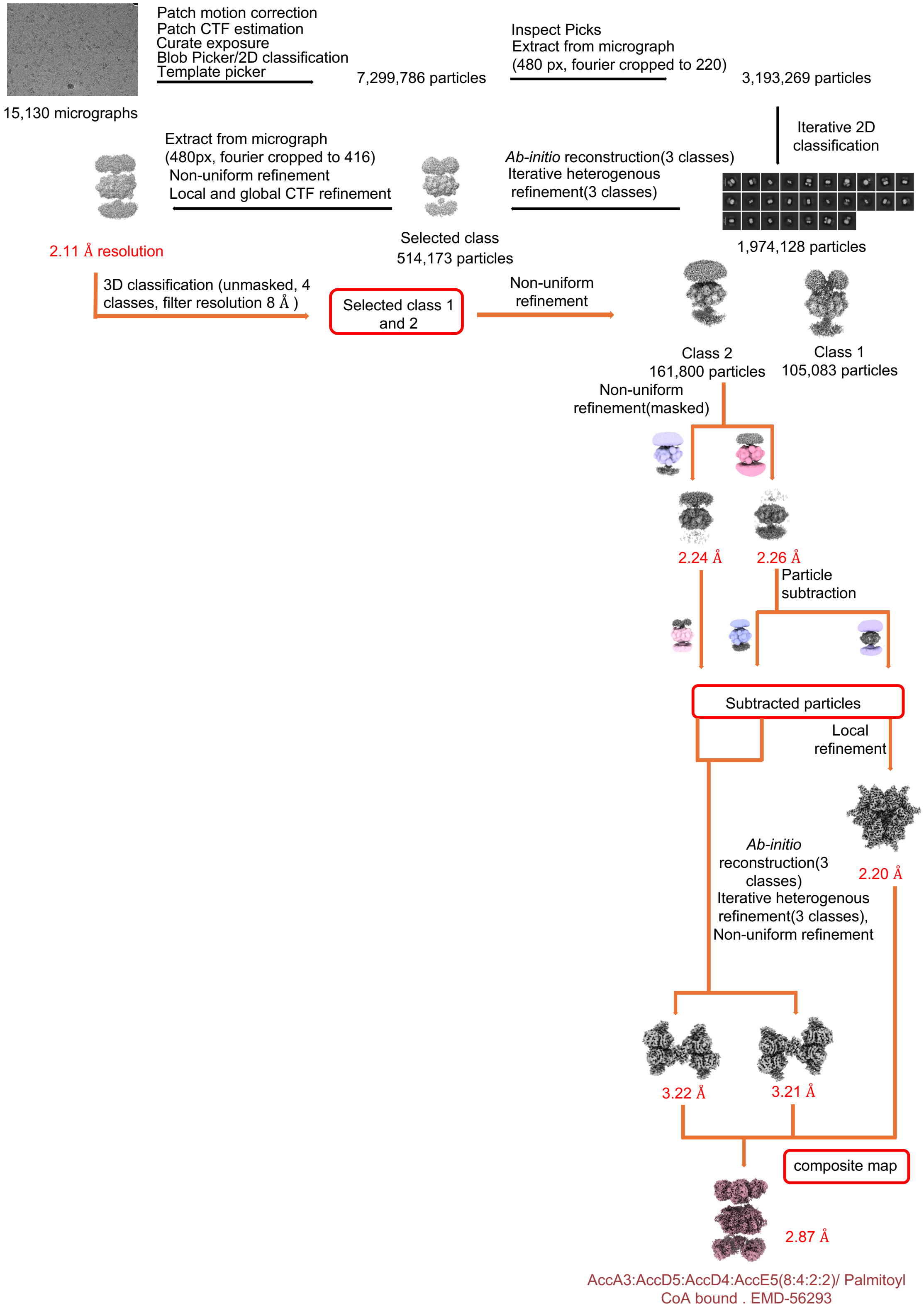


**Supplementary Fig. 7.** Image processing and classification workflow for the cryo-EM dataset of the LCC complex bound to palmitoyl-CoA, ADP, and $Mg^{2+}$.

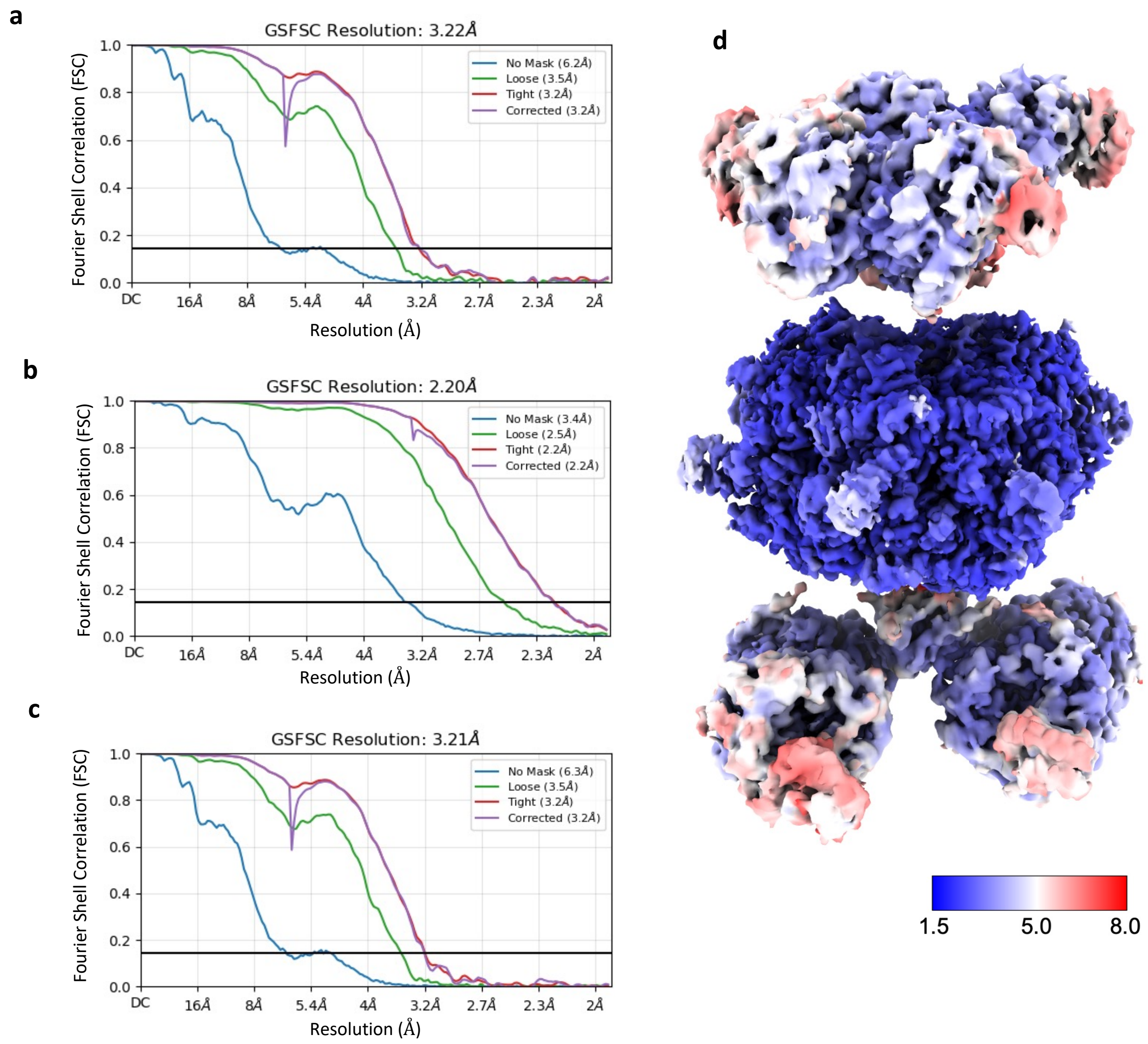


**Supplementary Fig. 8.** FSC curves for focused reconstructions of the palmitoyl-CoA–, $Mg^{2+}$–, and ADP-bound LCC complex: **a,** top BC module; **b,** central CT module; and **c,** bottom BC module. **d,** Estimated average resolution of the composite cryo-EM map.

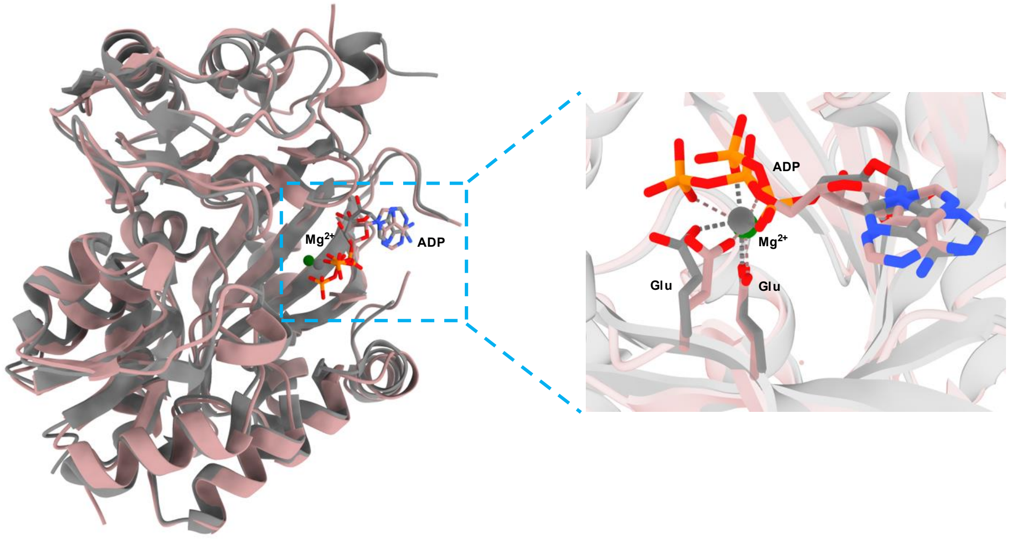


**Supplementary Fig. 9.** Structural superposition of AccA3 (brown) with the crystal structure of biotin carboxylase from Escherichia coli (PDB ID: 3RV3; grey), yielding an RMSD of 0.92 Å.

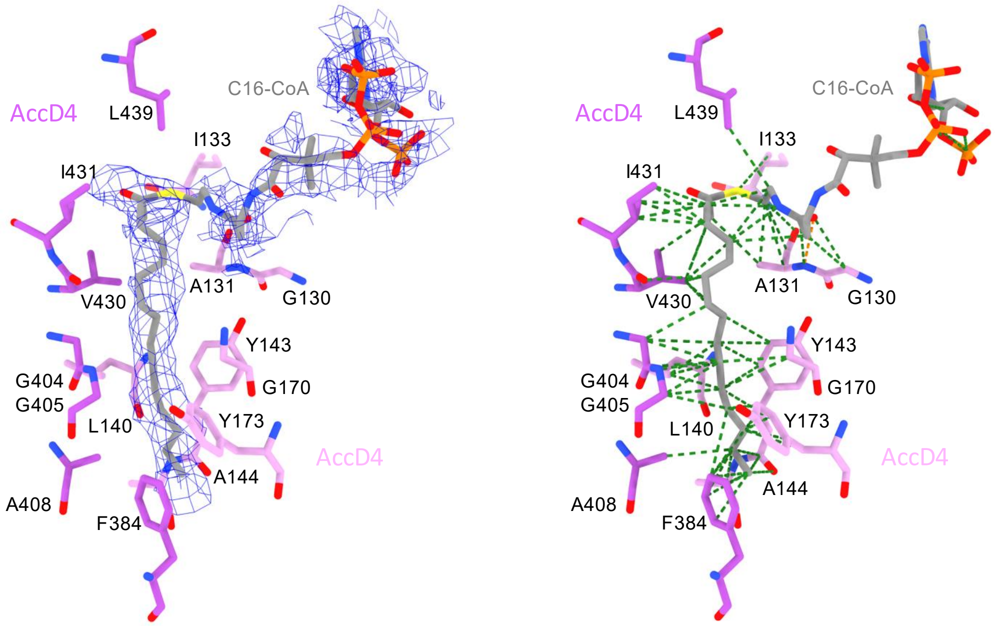


**Supplementary Fig. 10.** Under ATP-turnover conditions, no density corresponding to carboxylation at the C2 position of C16-CoA is observed. The acyl chain is well resolved (left) and primarily stabilized by van der Waals interactions (right, green dashed lines).

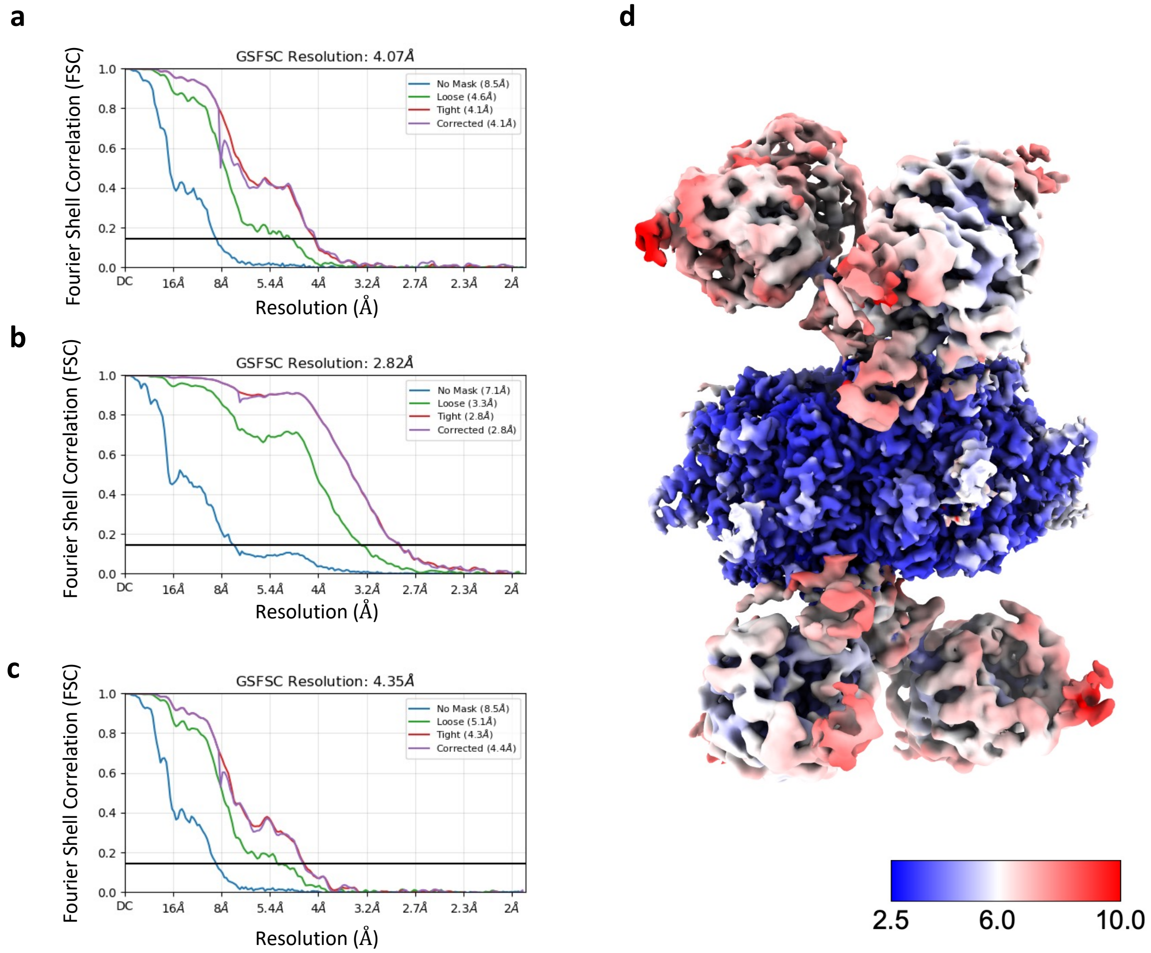


**Supplementary Fig. 11.** FSC curves for focused reconstructions of the propionyl-CoA– and palmitoyl-CoA–bound LCC complex: **a,** top BC module; **b,** central CT module; and **c,** bottom BC module. **d,** Estimated average resolution of the composite cryo-EM map.

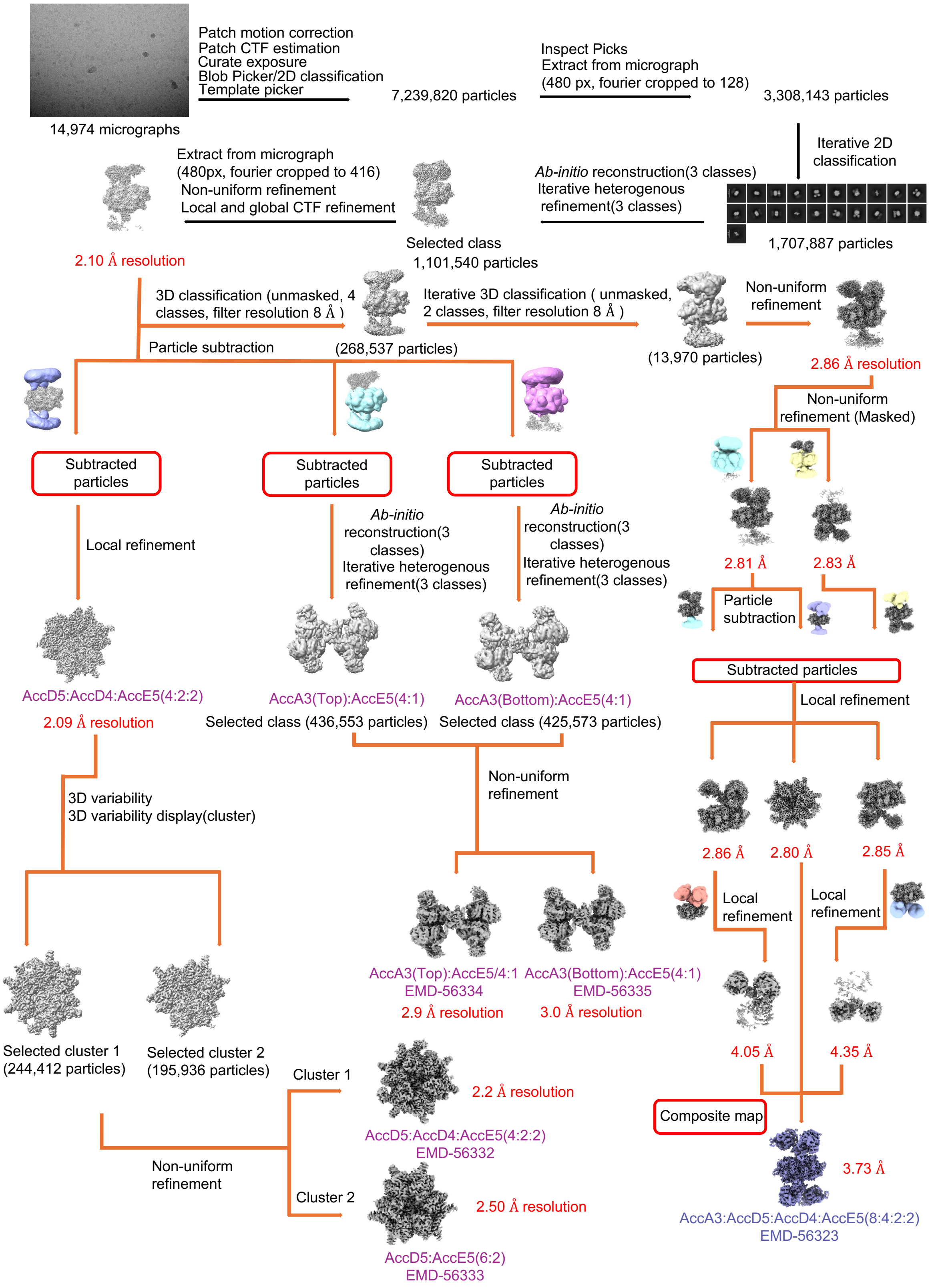


**Supplementary Fig. 12.** Image processing and classification workflow for the cryo-EM dataset of the LCC complex bound to propionyl-CoA and palmitoyl-CoA.

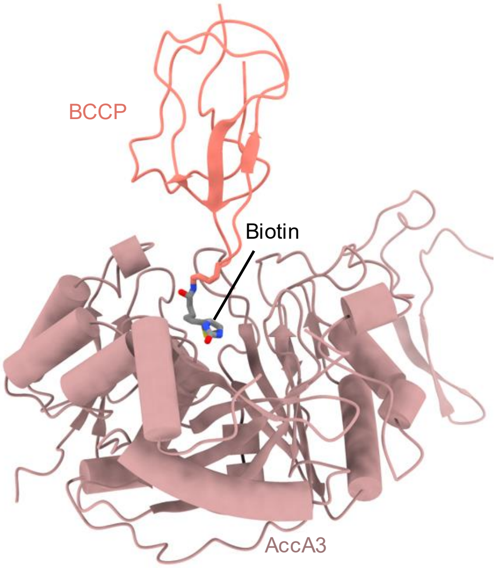


**Supplementary Fig. 13.** Biotinylated Lys564 oriented toward BC active site.

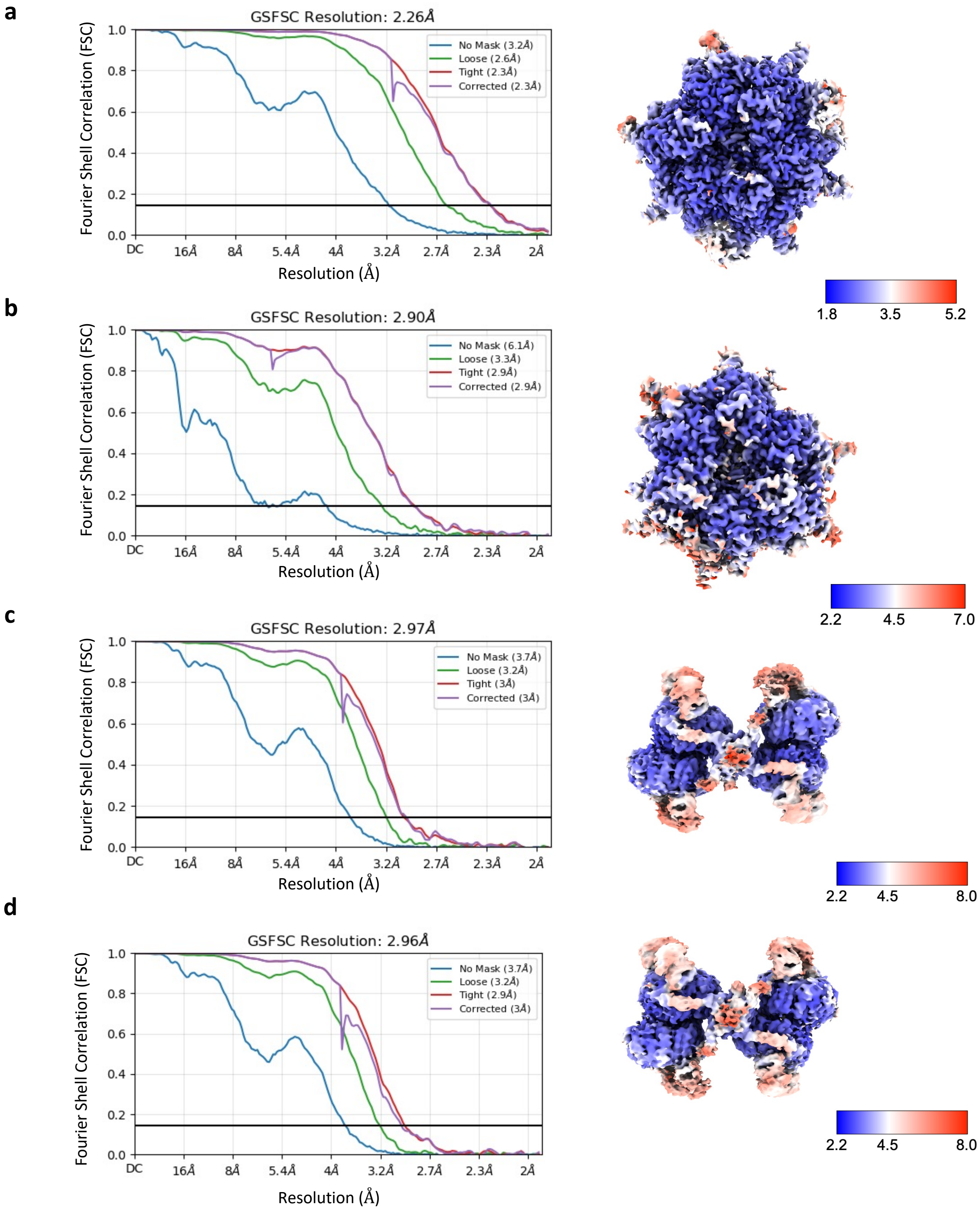


**Supplementary Fig. 14.** FSC curves (left) and local resolution estimates (right) for cryo-EM data of the palmitoyl-CoA–bound LCC complex: **a,** central CT module (4AccD5:2AccD4:2AccE5); **b,** central CT module (6AccD5:2AccE5); **c,** top BC module; and **d,** bottom BC module.

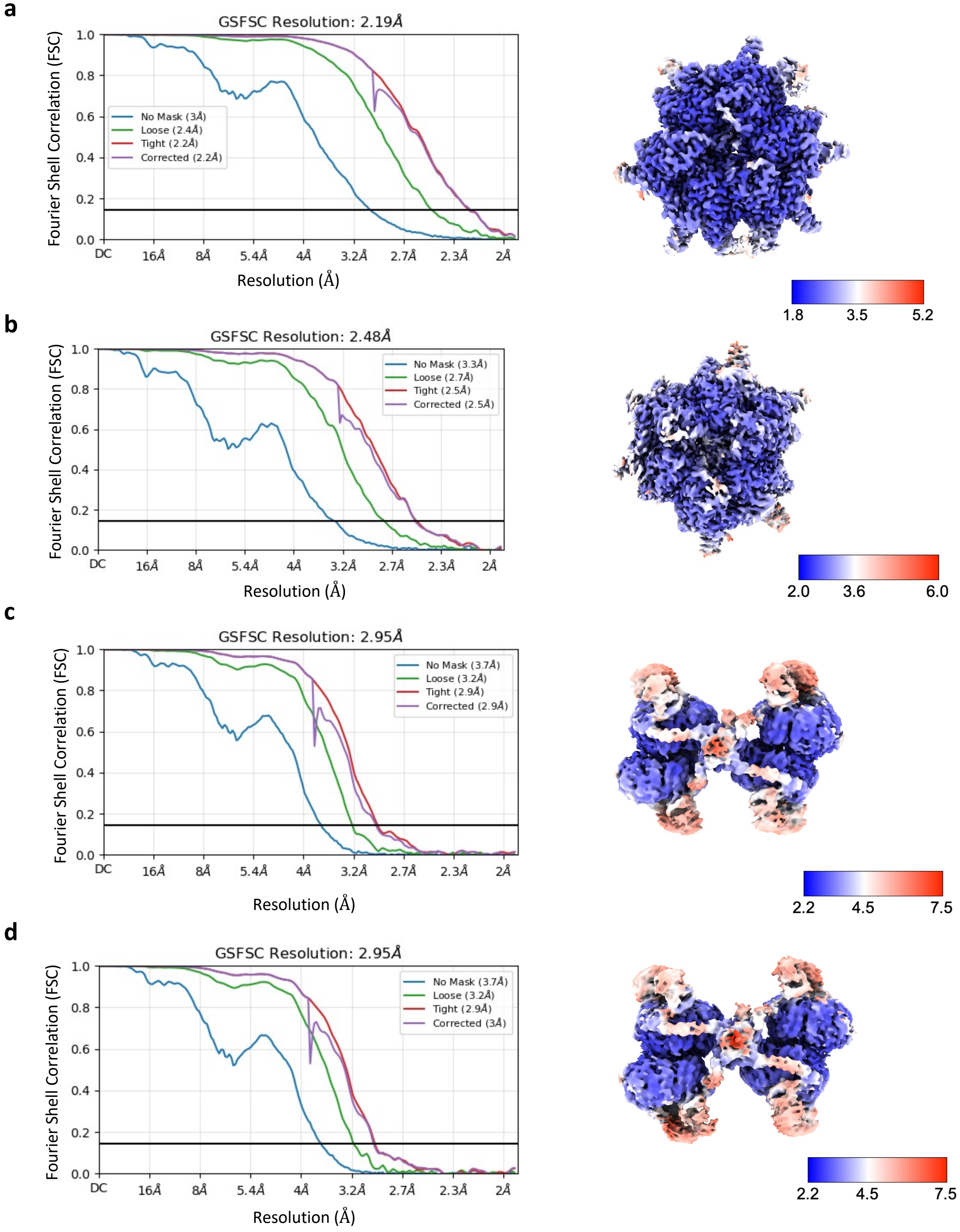


**Supplementary Fig. 15.** FSC plots (left) and local resolution estimation (right) from EM data of the LCC complex incubated with propionyl-CoA and palmitoyl-CoA. **a,** the central CT module (4AccD5:2AccD4:2AccE5), **b,** the central CT module (6AccD5:2AccE5), **c,** the top BC module, and **d,** the bottom BC module.

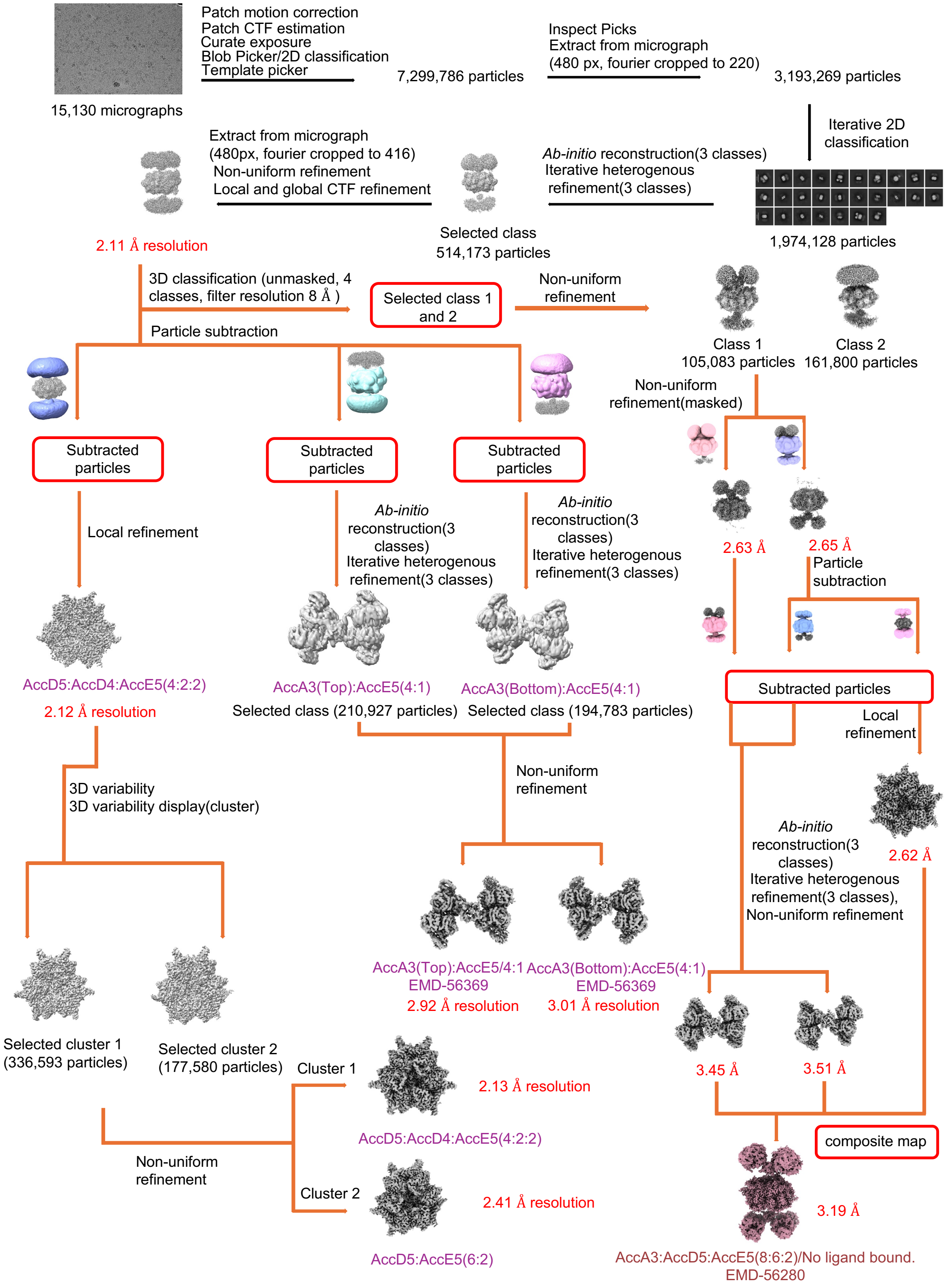


**Supplementary Fig. 16.** Image processing and classification workflow for the 8AccA3:6AccD5:2AccE5 complex bound to ADP and $Mg^{2+}$.

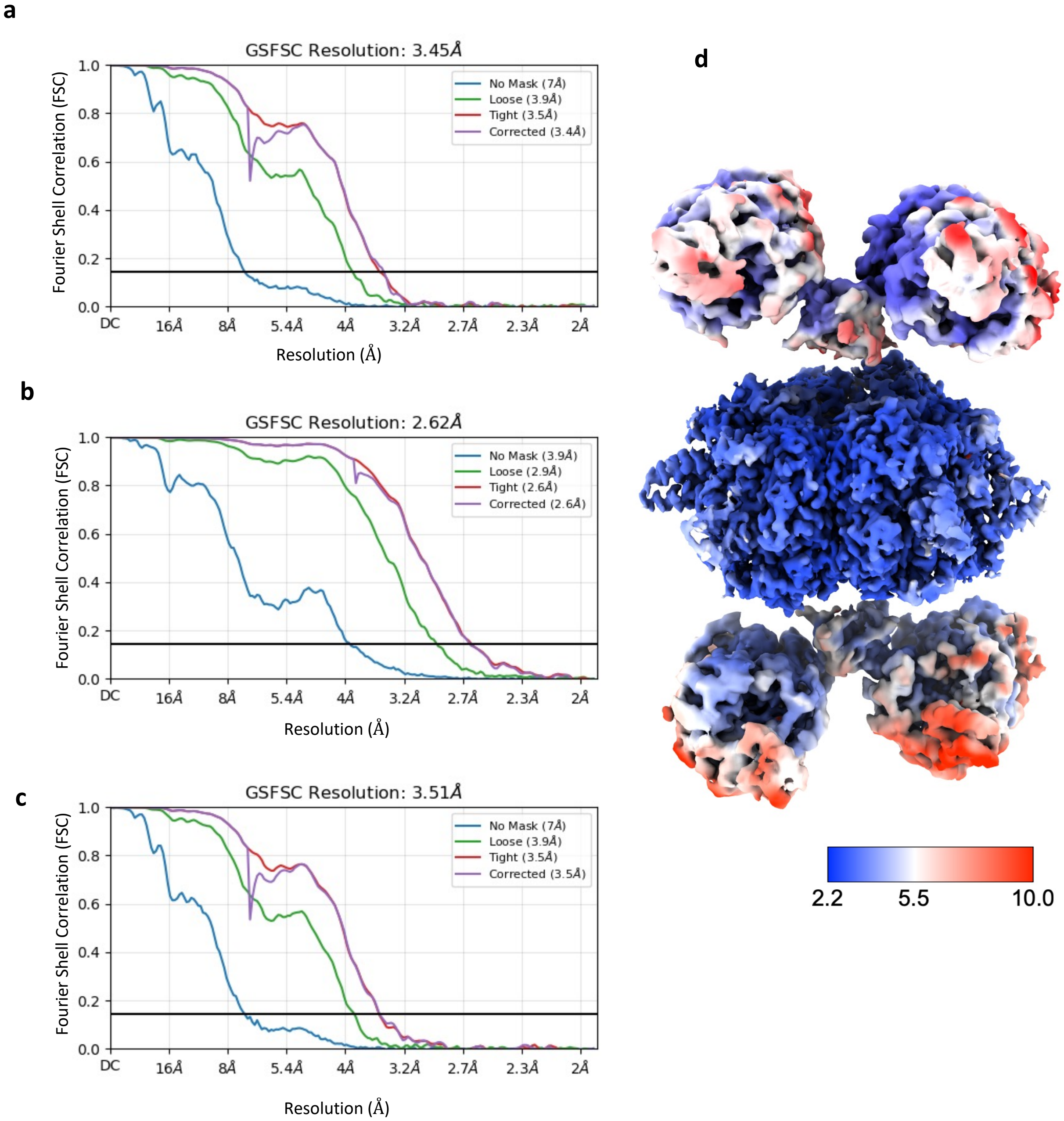


**Supplementary Fig. 17.** FSC curves for focused reconstructions of the $Mg^{2+}$– and ADP-bound 8AccA3:6AccD5;2AccE5 complex: **a,** top BC module; **b,** central CT module; and **c,** bottom BC module. **d,** Estimated average resolution of the composite cryo-EM map.

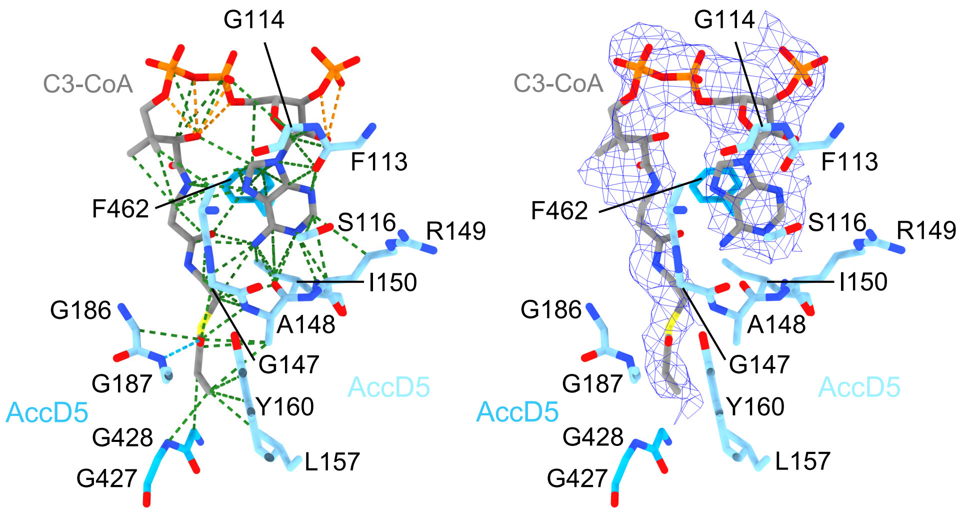


**Supplementary Fig. 18** Close-up view of C3-CoA interactions in the AccD5 active site. Hydrogen bonds (orange dashes) and van der Waals contacts (green dashes) are indicated. Right, cryo-EM density for C3-CoA (blue mesh).

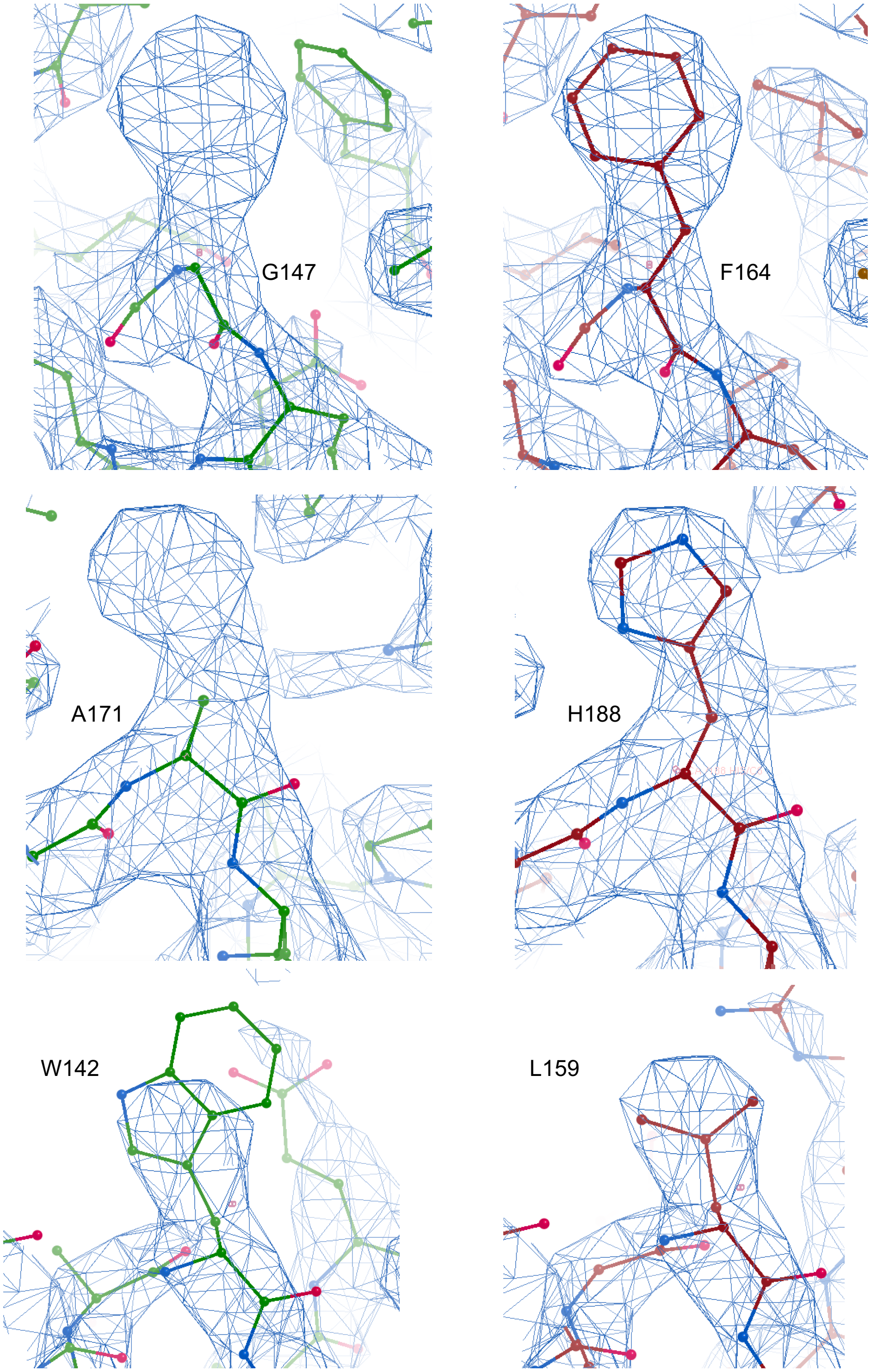


**Supplementary Fig. 19. Validation of the AccD5 homohexamericcore (ACCase 5).** Representative examples of model-map comparison at the carboxyltransferase active sites. Placement of AccD4-specific residues into the cryo-EM density (blue mesh; left) results in poor local model–map agreement, whereas substitution with the corresponding AccD5 residues (right) restores agreement with the density, confirming assignment of an AccD5 homohexameric core.

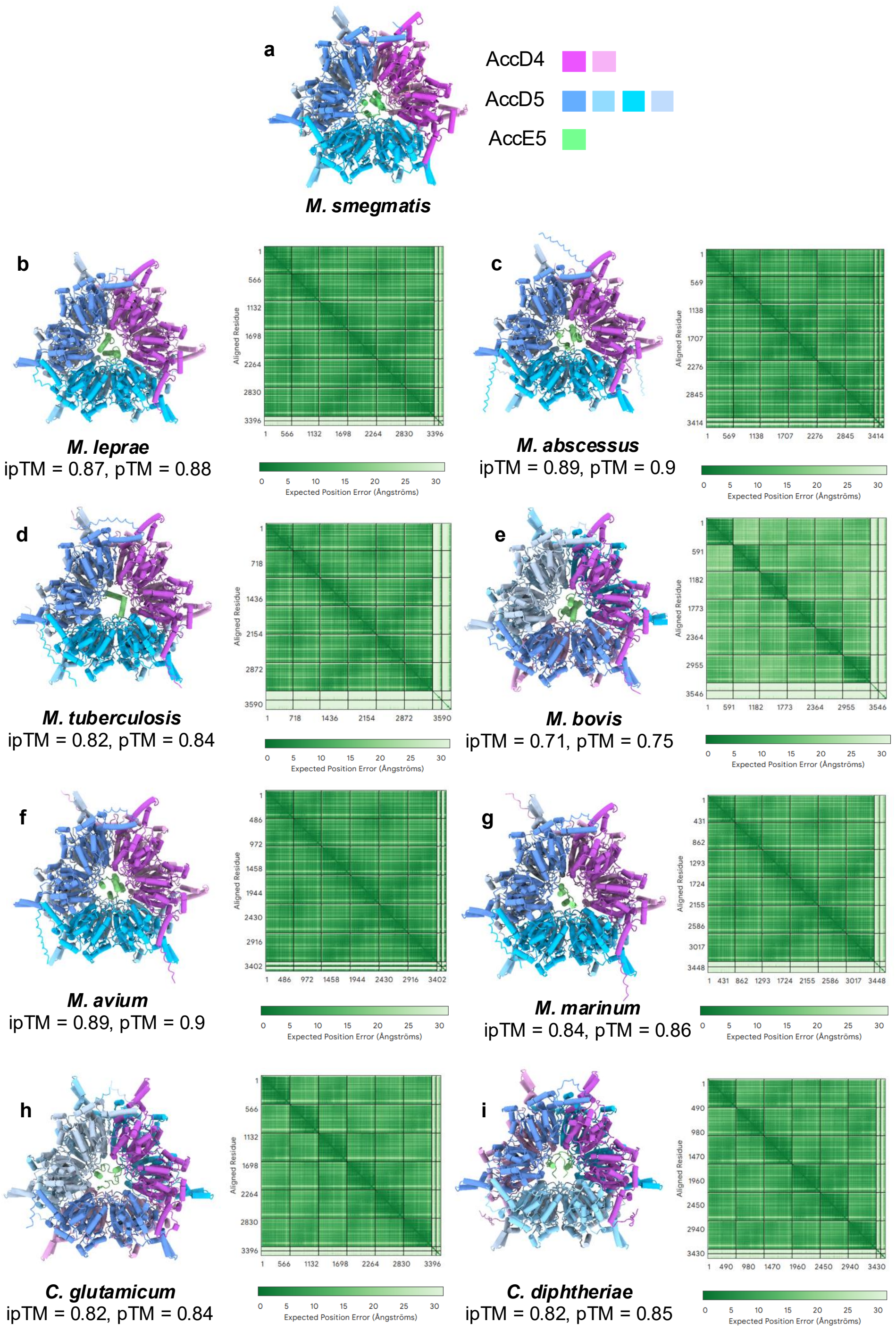


**Supplementary Fig. 20.** Predicted structures of the 4AccD5: 2AccD4::2AccE5 CT module across Corynebacteriales. The experimentally determined structure from Mycobacterium smegmatis **a** is compared with AlphaFold-predicted models from M. leprae **b**, M. abscessus **c**, M. tuberculosis **d**, M. bovis **e**, M. avium **f**, M. marinum **g**, Corynebacterium glutamicum **h**, and C. diphtheriae **i**.

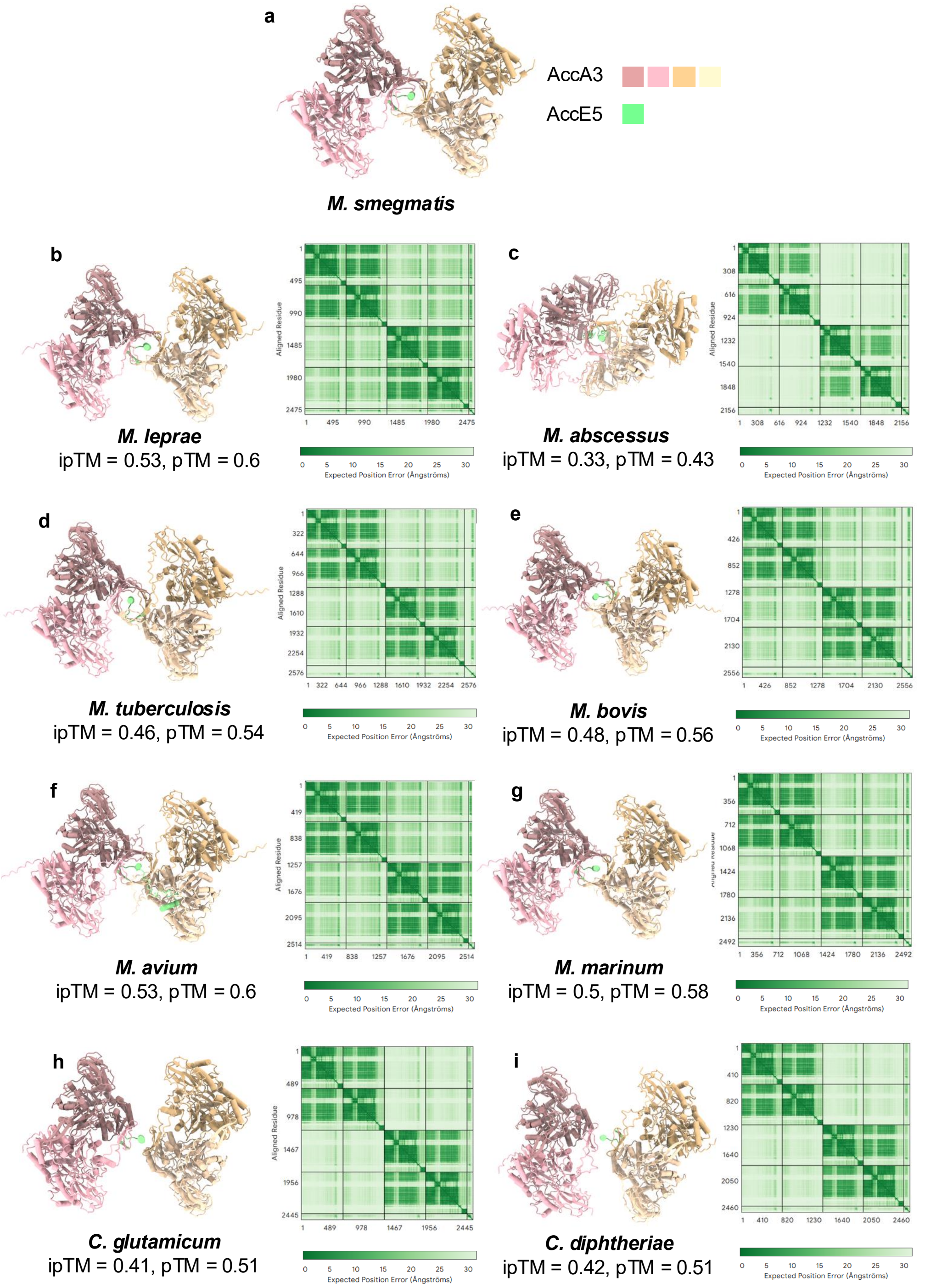


**Supplementary Fig. 21.** Predicted structures of the 4AccA3:1AccE5 BC module across Corynebacteriales. The experimentally determined structure from *Mycobacterium smegmatis* **a**, is compared with AlphaFold-predicted models from M. leprae **b,** *M. abscessus* **c**, *M. tuberculosis* **d**, *M. bovis* **e**, *M. avium* **f**, *M. marinum* **g**, *Corynebacterium glutamicum* **h**, and *C. diphtheriae* **i**.
